# Impaired peroxisomal import in *Drosophila* hepatocyte-like cells induces cardiac dysfunction through the pro-inflammatory cytokine Upd3

**DOI:** 10.1101/659128

**Authors:** Kerui Huang, Ting Miao, Kai Chang, Ping Kang, Qiuhan Jiang, Andrew J. Simmonds, Francesca Di Cara, Hua Bai

**Affiliations:** Department of Genetics, Development, and Cell Biology, Iowa State University, Ames, IA 50011, USA; Department of Cell Biology, University of Alberta, Edmonton, AB T6G 2H7, Canada.; Department of Microbiology & Immunology, Dalhousie University, Halifax, NS B3H4R2, Canada.

**Keywords:** Inflammaging, Cardiac aging, Oxidative stress, Peroxisomal import, Inter-organ communication, Oenocytes, Hepatocyte, Inflammatory cytokine, Interleukin-6.

## Abstract

Age is a major risk factor for cardiovascular diseases. Currently, the non-autonomous regulation of age-related cardiac dysfunction is poorly understood. In the present study, we discover that age-dependent induction of cytokine unpaired 3 (Upd3) in *Drosophila* oenocytes (hepatocyte-like cells), due to a dampened peroxisomal import function, is the primary non-autonomous mechanism for elevated arrhythmicity in old hearts. We show that Upd3 is significantly up-regulated (52-fold) in aged oenocytes. Oenocyte-specific knockdown of *Upd3* is sufficient to block aging-induced cardiac arrhythmia. We further show that the age-dependent induction of *Upd3* is triggered by impaired peroxisomal import and elevated JNK signaling in aged oenocytes. Intriguingly, oenocyte-specific over-expression of *Pex5*, the key peroxisomal import receptor, restores peroxisomal import, blocks age-related Upd3 induction, and alleviates aging- and paraquat-induced cardiac arrhythmicity. Thus, our studies identify an important role of the evolutionarily conserved pro-inflammatory cytokine signaling and hepatocyte-specific peroxisomal import in mediating non-autonomous regulation of cardiac aging.

## Introduction

Age is a major risk factor for a wide range of human diseases ^1^. For example, aging is associated with a great increase in the incidence of cardiovascular diseases (CVD), one of the leading causes of death worldwide ^2, 3^. During aging, cardiomyocytes undergo rapid remodeling with a variety of intracellular changes, in particular impaired mitochondrial quality control, increased production of reactive oxygen species (ROS), and elevated inflammation ^1^. The chronic and systemic inflammation (or “inflammaging”) is often associated with increased levels of circulating pro-inflammatory biomarkers (e.g., interleukin-6/IL-6 and C-reactive protein/CRP), which are notable risk factors for cardiovascular diseases ^4, 5^. The cause of inflammaging and its impact on cardiac aging are still poorly understood. It is known that short-term expression of inflammatory cytokines (e.g., IL-6) can protect myocytes from injury-induced apoptosis. However, prolonged production of IL-6 induces pathological hypertrophy and decreases cardiomyocyte contractility through the activation of Janus Kinases-Signal Transducer and Activator of Transcription (JAK-STAT) signaling ^6^. Elevated levels of circulating IL-6 are often associated with heart failure, myocardial damage and atherosclerosis ^6–8^. IL-6 can be produced not only by cardiomyocytes themselves in response to ischemia-reperfusion injury and myocardial infarction, but also by other neighboring tissues (e.g., endothelial cells and vascular smooth-muscle), immune cells (e.g., monocytes and macrophages), as well as liver ^7, 9^. However, the root causes of inflammaging and the primary sources of these inflammatory factors remain to be determined.

Liver is a major endocrine organ that produces a wide variety of systemic factors to coordinate body physiology and metabolism. It can produce pro-inflammatory cytokine IL-6 upon infection, injury, or partial hepatectomy ^10^. Patients with liver dysfunction, such as liver cirrhosis, often show increased cardiac arrhythmias, known as cirrhotic cardiomyopathy. Furthermore, it has been shown that nonalcoholic fatty liver disease (NAFLD) is a strong risk factor for cardiomyopathy ^11^. About 30% of alcoholic hepatitis patients develop cardiomyopathy and organ failure, suggesting a potential cross-talk between liver and heart. However, the direct link between liver dysfunction and cardiac aging remains elusive. It is known that aging significantly alters liver morphology and function, such as increased hepatocyte size, decreased mitochondrial number, altered lipid metabolism, and decreased expression of antioxidant enzymes such as superoxide dismutase 1 (*Sod1*) ^12^. Recently, using *Drosophila* oenocytes as a hepatocyte model, we observed a similar down-regulation of oxidative phosphorylation and detoxification, and up-regulation of inflammatory signaling in aged fly oenocytes ^13^. However, it remains unclear whether liver inflammation directly influences heart function at old ages.

Liver is known to enrich in peroxisome, a key organelle for ROS metabolism, alpha- and beta-oxidation of fatty acids, biosynthesis of ether phospholipid ^14^. The peroxisome assembly and the import of peroxisomal matrix proteins are controlled by a group of peroxisomal proteins called peroxins (PEXs). Mutations in peroxin genes disrupt normal peroxisome function and cause peroxisome biogenesis disorders (PBDs), such as Zellweger syndrome ^15^. Patients with Zellweger syndrome often develop severe hepatic dysfunction, such as hepatomegaly, cirrhosis, cholestasis, as well as inflammation ^16^. Several previous studies suggest that peroxisomal import function declines with age ^17–19^. Consistently, our recent translatomic analysis shows that the majority of peroxisome genes are down-regulated in aged fly oenocytes ^13^. In *Drosophila*, proper peroxisome function, in particular Pex5/Pex7-regulated peroxisome import, is required for phagocytosis and immune response in macrophages ^20^ and gut epithelium homeostasis ^21^. However, the role of peroxisome in aging regulation is currently unclear.

In this study, we discovered a peroxisome-mediated inter-organ communication between oenocytes and heart during *Drosophila* aging. We find that elevated ROS in aged oenocytes promotes cardiac arrhythmia non-autonomously through the pro-inflammatory factor unpaired 3 (Upd3), a four-helix bundle interleukin-6 (IL-6) type cytokine ^22^. Maintaining ROS homeostasis in oenocytes is sufficient to preserve cardiac function under aging or oxidative stress. Either reducing the expression of *Upd3* in oenocytes or blocking the activation of JAK-STAT signaling in cardiomyocytes alleviates aging- and oxidative stress-induced arrhythmia. Finally, we show that peroxisomal import function is significantly disrupted in aged oenocytes. *Pex5* knockdown-triggered peroxisomal import stress (PIS) up-regulates *Upd3* expression through c-Jun N-terminal Kinase (JNK) signaling in oenocytes. In contrast, oenocyte-specific over-expression of *Pex5* restores peroxisomal import, blocks age-induced Upd3 production and cardiac arrhythmicity. Together, our studies reveal a novel non-autonomous mechanism for cardiac aging that involves in hepatic peroxisomal import-mediated chronic and systemic inflammation.

## Results

### Oenocyte ROS homeostasis non-autonomously modulates cardiac function

Disrupted ROS homeostasis is one of the hallmarks of aging ^23^. Our recent translatomic analysis in *Drosophila* oenocytes (a hepatocyte-like tissue) revealed an overall down-regulation of anti-oxidant genes under aging, which is consistent with elevated oxidative stress in this tissue ^13^. To determine whether redox imbalance in oenocytes can non-autonomously impact cardiac function, we first induced oxidative stress specifically in oenocytes of female flies by crossing the *PromE-Gal4* driver ^24^ to RNAi lines against ROS scavenger genes *Catalase* (*Cat*) and *Superoxide dismutase 1* (*Sod1*) (Figure S1). Heart contractility was then assessed using the Semi-automatic Optical Heartbeat Analysis (SOHA) ^25^. Interestingly, oenocyte-specific knockdown (KD) of *Cat* and *Sod1* resulted in an increase in cardiac arrhythmicity, as measured by arrhythmia index (AI) (Figure 1a). These results suggest that disrupted ROS homeostasis in *Drosophila* oenocytes can modulate cardiac rhythm through an unknown non-autonomous mechanism.

**Figure 1:**
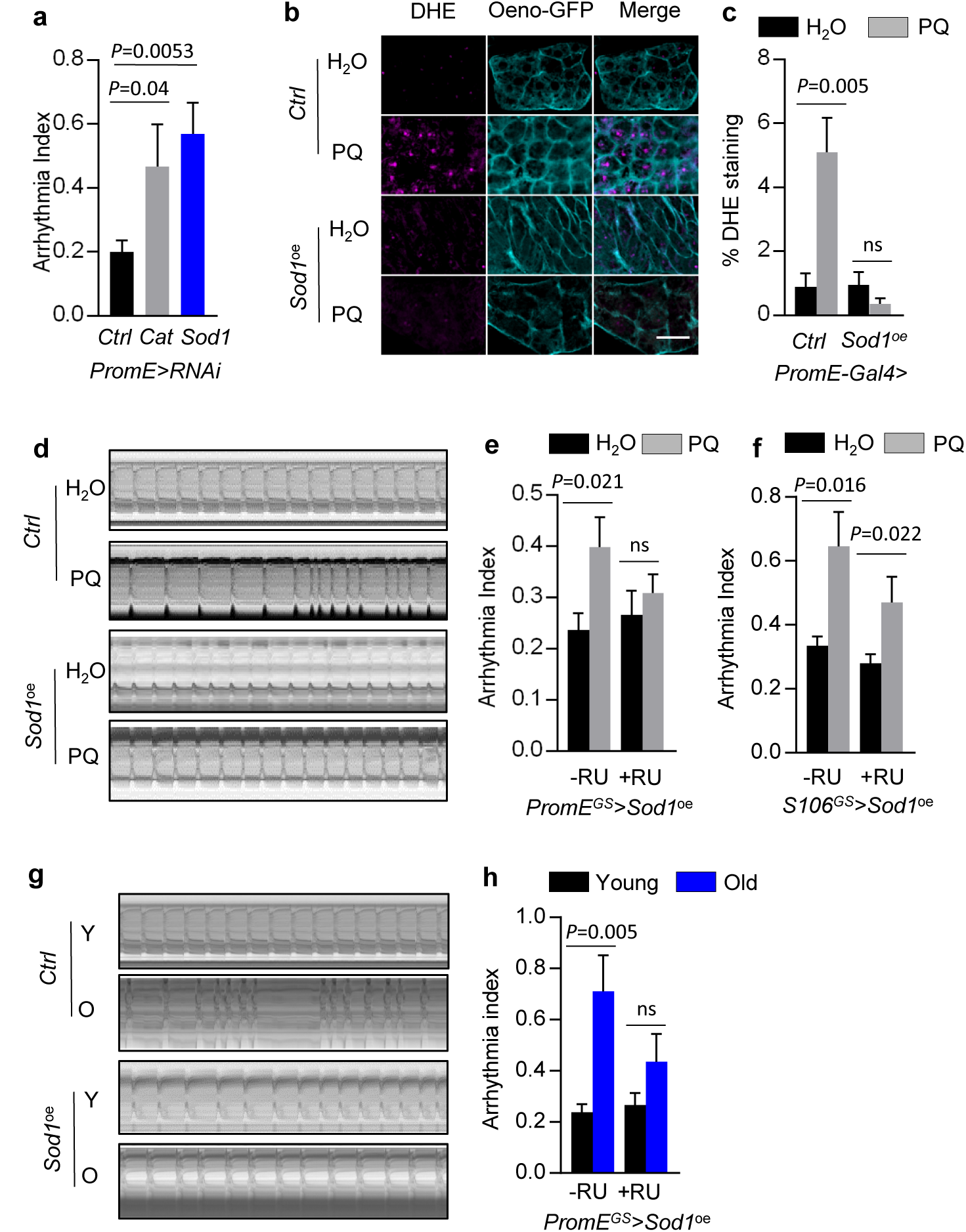
Oenocyte ROS homeostasis non-autonomously modulates cardiac function. **(a)** Arrhythmia index of oenocyte-specific *Cat* and *Sod1* knockdown flies (1-week-old). **(b)** Representative images of ROS levels (DHE staining) in dissected oenocytes from flies fed on normal diet or 10mM paraquat (for 24 h). All flies express mCD8::GFP under the oenocyte-specific driver (*PromE-Gal4*). *Sod1* was specifically over-expressed in the oenocytes (*Sod1*^oe^). Scale bar: 20 µm. **(c)** Quantification of the percentage of DHE-positive staining in a selected region of interest (n ≥ 16). **(d)** Representative M-mode traces showing heart contraction in control and *Sod1* over-expression flies fed on normal or 10mM paraquat diets. *Sod1* was expressed using the oenocyte-specific GeneSwitch driver (*PromE^GS^-Gal4*). RU: mifepristone (RU486). **(e)** Arrhythmia index of control and oenocyte-specific *Sod1* over-expression flies fed on normal or 10mM paraquat diets. **(f)** Arrhythmia index of control and fat body/gut-specific *Sod1* over-expression flies fed on normal or 10mM paraquat diets. over-expression specifically in fat body (*S106-Gal4*). **(g)** Representative M-mode traces showing heart contraction in young (2 weeks) and old (6 weeks) flies with or without oenocyte-specific *Sod1* over-expression. **(h)** Arrhythmia index of control and oenocyte-specific *Sod1*^oe^ flies at young and old ages. Data are represented as mean ± SEM. Panel e, f, h: N ≥ 16. Panel (a): N = 9. *P* values are calculated using unpaired *t-*test, ns: not significant.

Next, we asked whether heart function can be protected from oxidative stress and aging by maintaining redox balance in oenocytes. We first induced ROS level systemically by feeding flies with paraquat (PQ), an oxidative stress inducing agent. Feeding flies with paraquat for 24 hours induced ROS level in oenocytes, as measured by dihydroethidium (DHE) staining (Figures 1b-1c). Consistent with the previously report ^26^, paraquat feeding also induced arrhythmicity in fly hearts (Figures 1d-1e). Intriguingly, using an oenocyte-specific GeneSwitch driver (*PromE^GS^-Gal4*, Figure S2a), over-expression of *Sod1* in adult oenocytes (*PromE^GS^-Gal4>UAS-Sod1^oe^)* was sufficient to block PQ-induced ROS production in oenocytes (Figures 1b-1c), as well as alleviated PQ-induced arrhythmicity in the heart (Figures 1d-1e). Similarly, over-expressing *Sod1* in oenocytes attenuated aging-induced cardiac arrhythmicity (Figures 1g-1h). To examine whether Sod1-mediated cardiac protection is specific to oenocytes, we crossed *Sod1* over-expression line to a fat body/gut-specific GeneSwitch driver *S106^GS^-Gal4* ^27^ (Figure S2b). Over-expression of *Sod1* in fat body and gut did not rescue PQ-induced arrhythmia (Figure 1f). Together, these data suggest that oenocytes play a specific and crucial role in maintaining cardiac health during aging and PQ-induced oxidative stress, likely through an unknown circulating factor.

### Pro-inflammatory factor *Upd3* produced from oenocytes mediates aging- and paraquat-induced arrhythmia

To identify factors that are secreted by oenocytes and communicate to the heart to regulate cardiac function during aging and oxidative stress, we first compared the list of *Drosophila* secretory proteins ^28^ and our recent oenocyte translatomic analysis ^13^. We identified 266 secretory factors that are differentially expressed in aged or PQ-treated oenocytes (Figure 2a). Among these secretory factors, we selected 27 candidates that encode for cytokines and hormonal factors for a reverse genetic screen to determine their roles in mediating oenocyte-heart communication under oxidative stress. Knockdown of several candidate factors (e.g., Sala, BG642167) in oenocytes induced cardiac arrhythmia (Figure S3a), similar to the knockdown of *Cat* and *Sod1*. On the other hand, our genetic screening identified four candidates whose knockdown specifically in oenocytes significantly attenuated paraquat-induced cardiac arrhythmicity (Figure 2b). The four candidate genes are *PGRP-sb1*, *Ag5r2*, *TotA*, and *Upd3*. We further verified our screening results using oenocyte-specific GeneSwitch driver (*PromE^GS^-Gal4*) and repeated the knockdown experiments for *PGRP-sb1* (Figure S3b) and *Upd3* (Figure 2c, two independent *Upd3* RNAi lines used). The knockdown efficiency of *Upd3* RNAi was verified by QRT-PCR (Figure S4a). Consistent with the screening results, knockdown of *PGRP-sb1* and *Upd3* in adult oenocytes blocked paraquat-induced arrhythmia.

**Figure 2:**
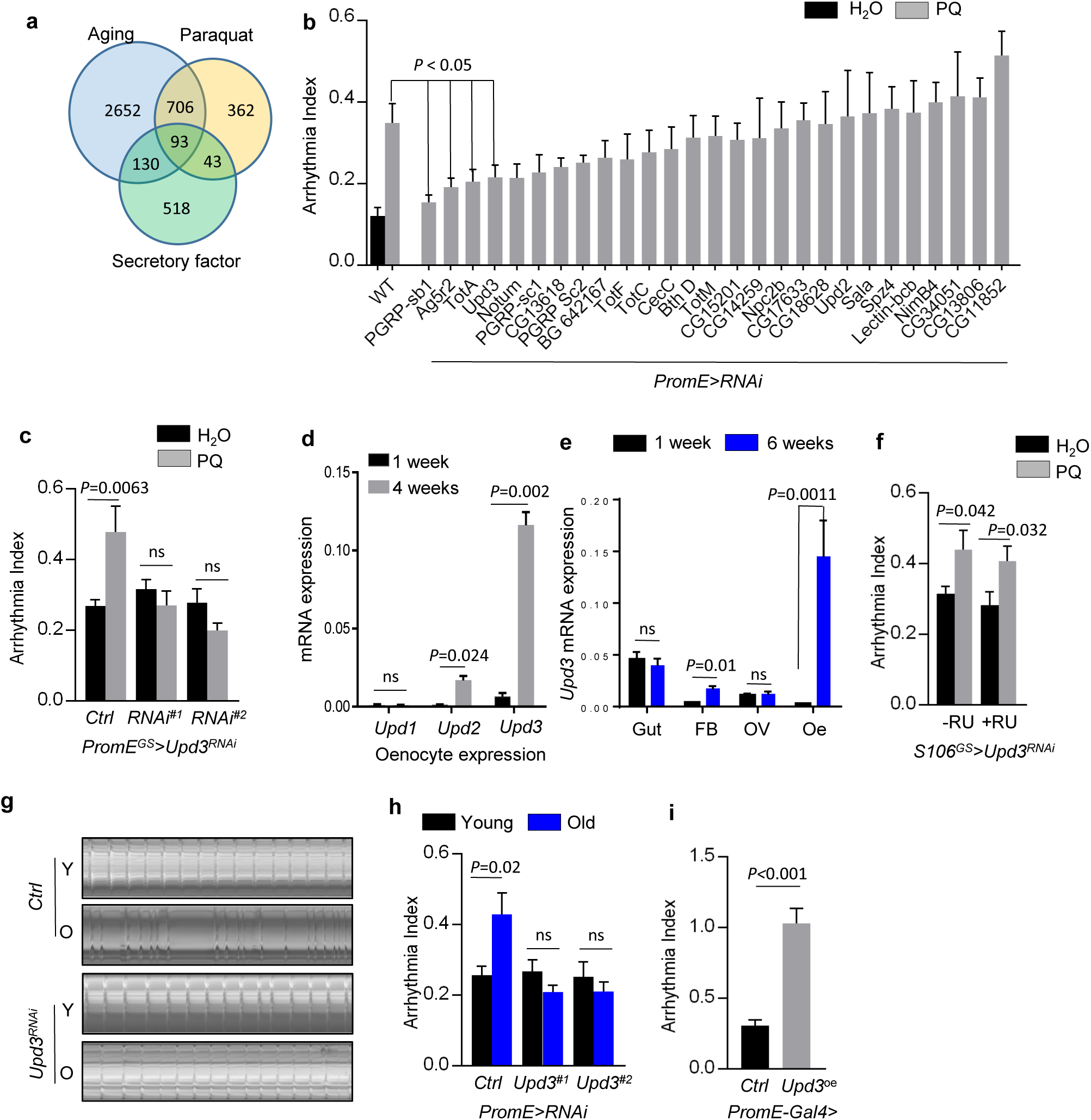
Pro-inflammatory factor *Upd3* produced from oenocytes mediates aging- and paraquat-induced arrhythmia. **(a)** Venn diagram showing the number of the predicted secretory proteins that are differentially expressed (≥ 2-fold, FDR<0.05) under aging and paraquat treatment. **(b)** Genetic screening on 27 candidate genes for their role in paraquat-induced arrhythmia. Fly are fed on 10 mM paraquat for 24 h before SOHA analysis. WT: Wild-type (*attP2* or *attP40* RNAi control lines), N ≥ 17. **(c)** Paraquat (PQ)-induced arrhythmia measured by SOHA for two independent *Upd3* RNAi lines under oenocyte-specific GeneSwitch driver (*PromE^GS^-Gal4*). N ≥ 18. **(d)** Relative mRNA expression of *Upd1*, *Upd2* and *Upd3* from isolated oenocytes at ages of 1 week or 4 weeks. N = 3. **(e)** Relative mRNA expression of *Upd3* in different tissues dissected from young (1 week) and old (6 weeks) female flies. FB: fat body, OV: ovary, oe: oenocytes. N = 3. **(f)** PQ-induced arrhythmia measured by SOHA for *Upd3 RNAi* under fat body-specific GeneSwitch driver (*S106^GS^-Gal4*). N ≥ 17. **(g)** Representative M-mode traces of wild-type and oenocyte-specific *Upd3* knockdown flies at young and old ages. **(h)** Arrhythmia index of wild-type and oenocyte-specific *Upd3* knockdown flies. Two independent RNAi lines used. N ≥ 17. **(i)** Arrhythmia index for flies with ectopic *Upd3* expression (*UAS-Upd3-GFP*) specifically in oenocytes. N ≥ 17. Data are represented as mean ± SEM. *P* values are calculated using unpaired *t-*test, ns: not significant.

Among the identified secretory factors, *Upd3* is a pro-inflammatory factor that belongs to the four-helix bundle interleukin-6 (IL-6) type cytokine family ^22^. In *Drosophila*, *Upd3* is one of the three ligands that activate the JAK/STAT signaling pathway. The expression of *Upd3* in oenocytes was higher than two other unpaired proteins (*Upd1* and *Upd2*), and *Upd3* expression was significantly induced under aging (Figure 2d). Although the expression of *Upd2* was slightly induced in aged oenocytes (Figure 2d), *Upd2* KD did not block paraquat-induced arrhythmicity (Figure 2b). Furthermore, we found that *Upd3* transcripts can be detected in several other adult tissues besides oenocytes, such as abdominal fat body, gut and ovary (Figure 2e). The highest expression of *Upd3* was found in the gut at young ages (Figure 2e). Intriguingly, *Upd3* expression in oenocytes increased sharply during normal aging (more than 52-fold) and became the highest among all adult tissues at old ages. *Upd3* expression did not show age-dependent increases in gut and ovary, and it only slightly increased in aged fat body (Figure 2e). These findings suggest that oenocytes are the primary source of *Upd3* production in aged flies.

Because *Upd3* expressed in multiple adult tissues, we wonder if Upd3 produced from tissues other than oenocytes also contributes to the non-autonomous regulation of cardiac function. To test this tissue-specific effect, we knocked down *Upd3* using the fat body/gut-specific driver (*S106^GS^-Gal4*) and found that fat body/gut-specific *Upd3* KD did not alleviate paraquat-induced arrhythmia (Figure 2f). Similar to the findings from paraquat treatment (Figure 2c), oenocyte-specific *Upd3* KD also blocked aging-induced cardiac arrhythmia (Figures 2g-2h, two independent *Upd3* RNAi lines showed). Conversely, oenocyte-specific over-expression of *Upd3* at young ages induced premature cardiac aging phenotypes (high arrhythmia index) (Figure 2i), which is similar to age-induced cardiac arrhythmia seen in multiple control flies (Figure S3c). Taken together, these results suggest that *Upd3* is the primary cytokine that is secreted from oenocytes to regulate aging- and stress-induced cardiac arrhythmicity.

### Oenocyte-produced *Upd3* activates JAK-STAT pathway in cardiomyocytes to induce arrhythmia

*Upd3* is known to systemically up-regulation JAK-STAT pathway in response to tissue injuries, excess dietary lipid, and ingestion of paraquat ^29–31^. We asked whether oenocyte-produced *Upd3* can signal to the heart and activate JAK-STAT pathway in cardiomyocytes. To test this idea, we used the nuclear localization of STAT (also known as *Stat92E* in *Drosophila*) to indicate the activation of JAK-STAT signaling. Consistent with previous finding ^31, 32^, paraquat treatment induced the levels of transcription factor STAT and promoted its localization to the nucleus in heart tissue (Figures 3a-3b). Interestingly, we found that oenocyte-specific *Upd3* KD attenuated paraquat-induced STAT nuclear localization in the heart (Figures 3a-3b).

**Figure 3:**
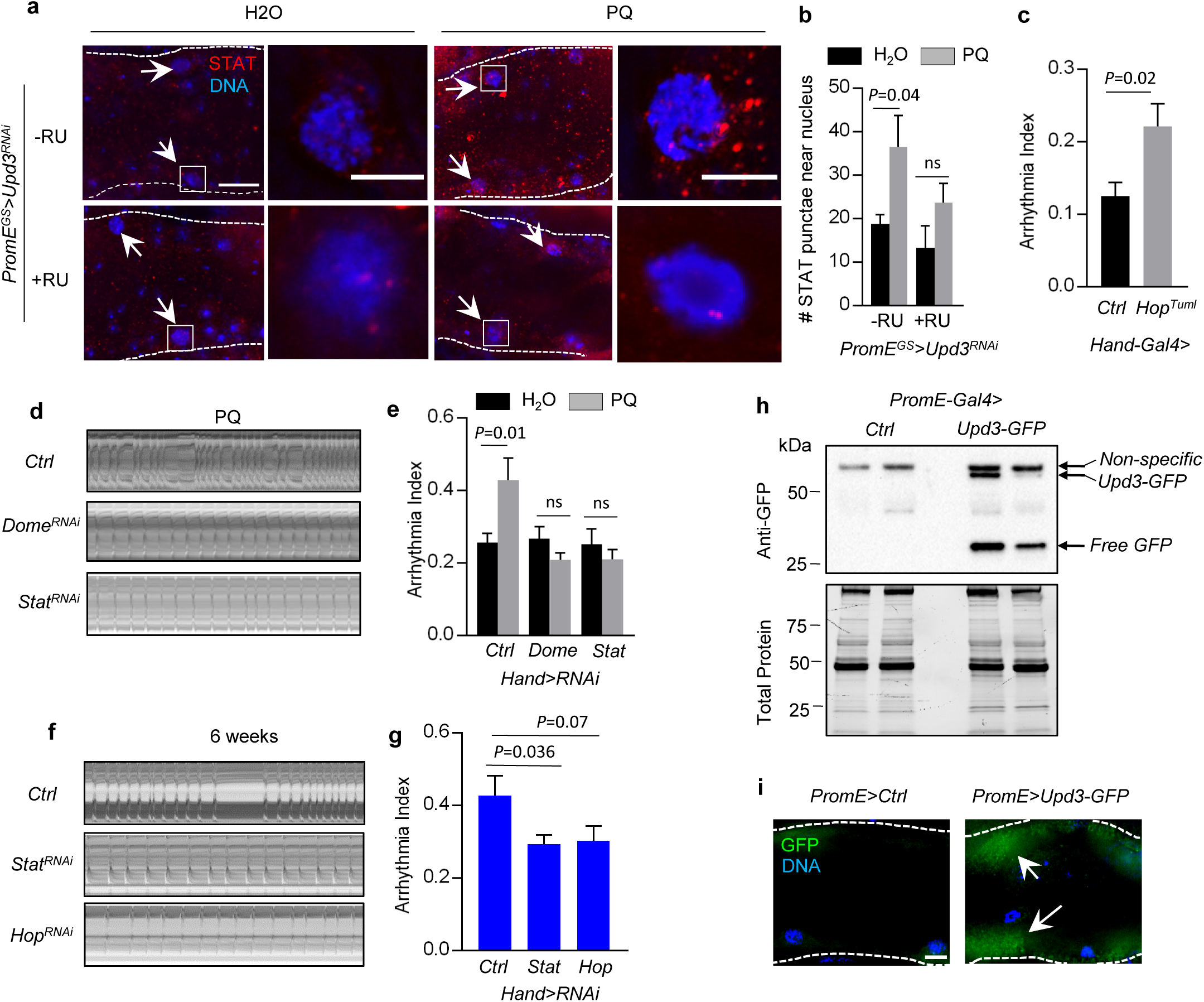
Oenocyte-produced *Upd3* activates JAK-STAT pathway in cardiomyocytes to induce arrhythmia. **(a)** Representative images of STAT immunostaining in the heart of oenocyte-specific *Upd3* KD (+RU) and control flies (-RU). Flies were treated with normal or paraquat diet for 24 h. Arrows indicate cardiomyocyte nuclei. White boxes indicate the regions shown in the right insets. Scale bar: 20 µm (inset: 5 µm). **(b)** Quantification of the STAT-positive punctae near cardiomyocyte nucleus. N = 7. **(c)** Arrhythmia index of young flies (1-week-old) with heart-specific expression of an activated form of *Hop* (*Hop^Tuml^*). N ≥ 21. **(d)** Representative M-mode traces of paraquat-treated wild-type, heart-specific *Dome* and *Stat* RNAi flies. **(e)** Arrhythmia Index of paraquat-treated wild-type, heart-specific *Dome* and *Stat* KD flies. N ≥ 15. **(f)** Representative M-mode traces of heart-specific *Stat* and *Hop* RNAi flies at old ages. **(g)** Arrhythmia Index of heart-specific *Stat* and *Hop* RNAi flies at old ages. N ≥ 18. **(h)** Western blot analysis on the hemolymph samples extracted from flies expressing Upd3-GFP fusion proteins specifically in oenocytes. Two biological replicates are shown. Total protein loaded onto the Bio-Rad Stain-Free gel was visualized using ChemiDoc MP Imagers after UV activation. **(i)** Immunostaining of heart tissues using anti-GFP antibody from flies expressing *Upd3-GFP* in oenocytes. Scale bar: 20 µm. Data are represented as mean ± SEM. *P* values are calculated using unpaired *t-*test, ns: not significant.

Activation of JAK-STAT play a significant role in the pathogenesis of myocardial ischemia and cardiac hypertrophy ^33, 34^. We then asked whether blocking JAK-STAT signaling in fly hearts could protect cardiac function under oxidative stress and aging. As expected, we found that heart-specific activation of JAK-STAT signaling by expressing an active form of JAK kinase *Hopscotch/Hop* (*Hop^Tuml^)* in young fly heart (using heart-specific driver *Hand-gal4*) induced cardiac arrhythmia prematurely (Figure 3c). Conversely, cardiac-specific knockdown of either the receptor *Domeless (Dome)* or the transcription factor *Stat* blocked PQ-induced cardiac arrhythmia (Figures 3d-3e). Similarly, knockdown of *Stat* and *Hop* in the heart attenuated aging-induced arrhythmia (Figures 3f-3g).

Next, we asked whether oenocyte-produced *Upd3* is secreted into the hemolymph and directly targeted cardiomyocytes. We over-expressed Upd3-GFP fusion proteins specifically in the oenocytes and analyzed the hemolymph samples using western blotting. The Upd3-GFP fusion proteins were successfully detected in the hemolymph extracted from *PromE* >*Upd3-GFP* flies (Figure 3h), suggesting the oenocyte-produced Upd3 indeed can be secreted into the hemolymph. Interesting, free GFP proteins were also found in the hemolymph, which may be due to a cleavage of the C-terminus of Upd3 occurring after its secretion (Figure 3h).

Consistently, we found high levels of GFP signals in the heart tissue dissected from the flies expressing *Upd3-GFP* under the oenocyte-specific driver (*PromE-Gal4*) (Figure 3i). Together, these data suggest that Upd3 produced from oenocytes is released into the hemolymph and activate JAK-STAT in the heart to regulate cardiac function.

### Impaired peroxisomal import in oenocytes induces *Upd3* expression and promotes cardiac arrhythmia

Next, we asked how aging up-regulates *Upd3* expression in oenocytes. In our previous oenocyte translatomic analysis, we found that genes involved in oxidative phosphorylation and peroxisome biogenesis are significantly down-regulated during aging, which is consistent with elevated ROS levels in aged oenocytes ^13^. It is known that mitochondria and peroxisome are two major ROS contributors. We then investigated whether age-dependent down-regulation of genes in mitochondrial respiratory chain complexes and peroxisome biogenesis contributes to Upd3 over-production and oenocyte-heart communication. Interestingly, oenocyte-specific knockdown of mitochondrial complex I core subunit *ND75* and mitochondrial manganese superoxide dismutase *Sod2* ^35, 36^ showed no effects on cardiac arrhythmia (Figure 4b). The knockdown efficiency of *ND75* RNAi was verified by QRT-PCR (Figure S4b). On the other hand, knockdown of the key factors involved in peroxisomal import process (*Pex5, Pex1 and Pex14*) in oenocytes significantly induced cardiac arrhythmia (Figure 4b). The knockdown efficiency of *Pex1* RNAi and *Pex5* RNAi was verified by QRT-PCR (Figure S4c-S4d). Pex5 is the key import factor that binds to cargo proteins containing peroxisomal targeting signal type 1 (PTS1) and delivers them to peroxisomal matrix through Pex13/Pex14 docking complex and Pex1/Pex6 recycling complex ^37, 38^ (Figure 4a). Interestingly, Pex5 itself is the major component of the peroxisomal translocon (also known as importomer, or import pore), which interacts with Pex14 and translocates cargo proteins across peroxisomal membrane through an ATP-independent process ^39–41^. In addition, we noticed that not all peroxisome genes were involved in oenocyte-heart communication. Oenocyte-specific knockdown of *Pex19*, the key peroxisomal membrane assembly factor, did not promote cardiac arrhythmia (Figures 4a-4b).

**Figure 4.**
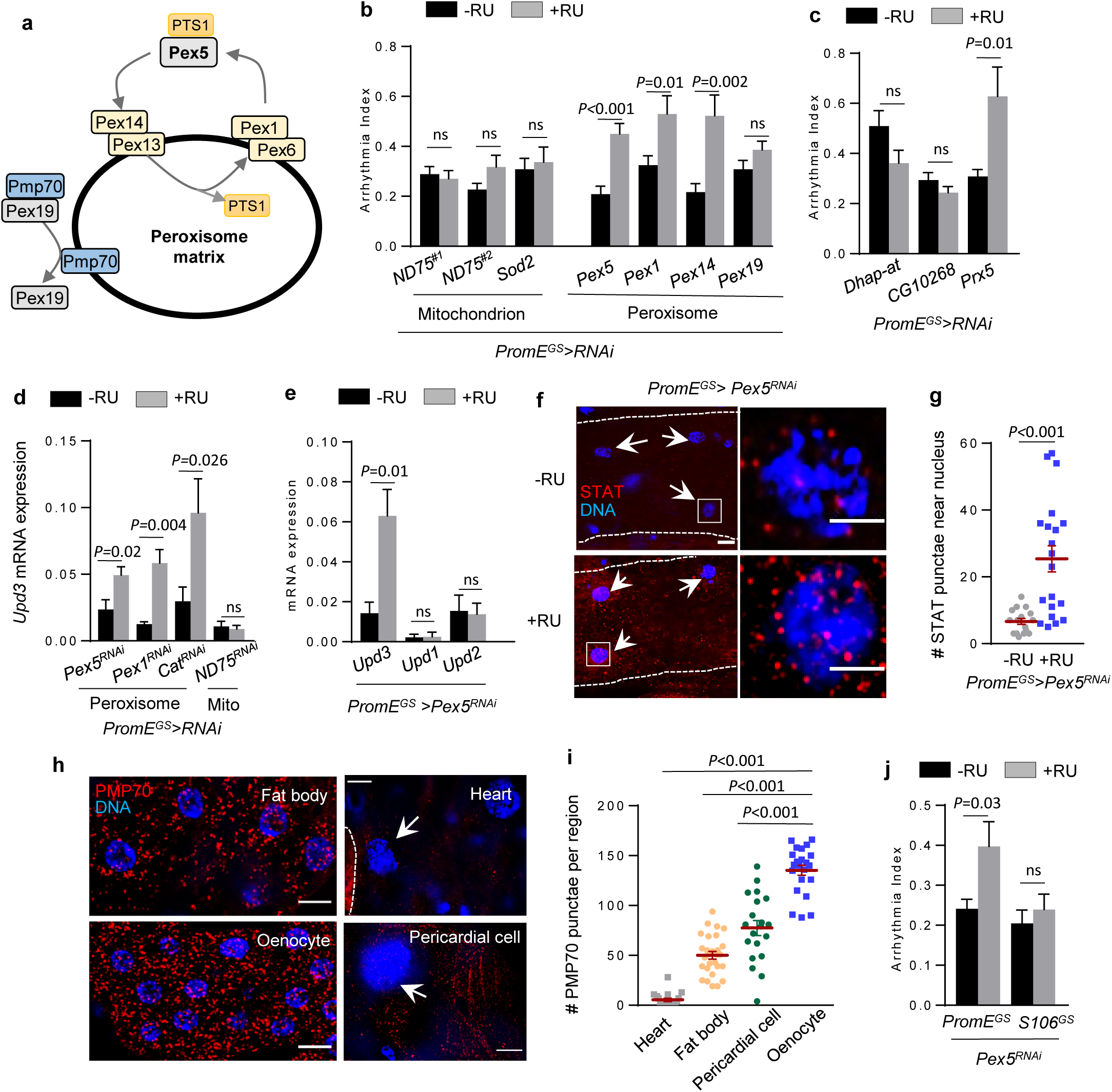
Impaired peroxisomal import in oenocytes induces *Upd3* expression and promotes cardiac arrhythmia. **(a)** Schematic diagram showing the key genes involved in peroxisomal import and membrane assembly. **(b)** Arrhythmia index of oenocyte-specific knockdown of mitochondrial complex I subunit *ND75* (two independent RNAi lines), mitochondrial Mn superoxide dismutase *Sod2*, peroxisomal import factors (*Pex5, Pex1*, *Pex14*), and peroxisomal membrane assembly factor (*Pex19*). N ≥ 15. Oenocyte-specific GeneSwitch driver (*PromE^GS^-Gal4*) was used. RU: mifepristone (RU486). **(c)** Arrhythmia index of oenocyte-specific knockdown of peroxisomal matrix enzymes, *Dhap-at*, *CG10268*, *Prx5*. N ≥ 15. **(d)** QRT-PCR analysis showing relative mRNA levels of *Upd3* from dissected oenocytes of *Pex5*, *Pex1*, *Cat* and *ND75* KD flies. Mito: Mitochondrion. N = 4∼6. **(e)** QRT-PCR analysis showing relative mRNA levels of *Upd3*, *Upd1*, *Upd2* from dissected oenocytes of oenocyte-specific *Pex5* KD flies. N = 4∼6. **(f)** Representative images of STAT immunostaining of heart tissues dissected from flies with (+RU) or without (-RU) oenocyte-specific *Pex5* KD. Arrows indicate cardiomyocyte nuclei. White boxes indicate the regions shown in the right insets. Scale bar: 20 µm (inset: 5 µm). **(g)** Quantification on the number of STAT punctae around cardiomyocyte nuclei of oenocyte-specific *Pex5* KD flies. N = 6. Dot plot shows the quantifications of 6 biological replicates, 3-4 selected regions of interest (ROIs) per replicate. **(h)** Representative images of PMP70 immunostaining in three fly tissues. Arrows indicate cardiomyocyte and pericardial cell nuclei. Scale bar: 6.7 µm. **(i)** Quantification of PMP70-positive peroxisomes per region of interest. N = 6. Dot plot shows the quantifications of 6 biological replicates, 3-5 ROIs per replicate. **(j)** Arrhythmia index of flies with *Pex5* KD in either oenocytes (*PromE^GS^-Gal4*) or fat body/gut (*S106^GS^-Gal4*). N ≥ 15. Data are represented as mean ± SEM. *P* values are calculated using unpaired *t-*test, ns: not significant.

Our previous oenocyte translatomic analysis found that many peroxisomal matrix enzymes were significantly down-regulated during aging, such as peroxiredoxin-5 (*Prx5*, a thioredoxin peroxidase regulating hydrogen peroxide levels), dihydroxyacetone phosphate acyltransferase (*Dhap-at*, the key enzyme catalyzing the first step in the biosynthesis of ether phospholipid, such as plasmalogens), and phosphomevalonate kinase (*CG10268*, the enzyme involved in mevalonate pathway). Interestingly, knockdown of *Prx5*, but not *Dhap-at* and *CG10268*, induced cardiac arrhythmia (Figure 4c). Thus, these data suggest that peroxisome-mediated ROS homeostasis, rather than lipid metabolism, plays an important role in modulating cardiac function.

We then asked whether impaired peroxisome import in oenocytes influences *Upd3* expression. As expected, knockdown of *Pex5* or *Pex1* in oenocytes induced *Upd3* mRNA levels (Figure 4d). Similarly, knockdown of peroxisomal antioxidant gene *Cat* significantly up-regulated *Upd3* transcription, whereas inhibition of mitochondrial complex I subunit *ND75* did not altered *Upd3* expression (Figure 4d). Interestingly, the other two unpaired proteins *Upd1* and *Upd2* remained unchanged under *Pex5* KD (Figure 4e). Given that impaired peroxisome function induces *Upd3* expression in oenocytes, we speculate that *Pex5* KD in oenocytes could remotely regulate JAK-STAT activity in the heart. Indeed, we found that oenocyte-specific *Pex5* KD induced STAT expression and STAT nuclear localization in the heart (Figures 4f-4g). Together, we show that impaired peroxisomal import promotes Upd3 production in oenocytes and activate cardiac JAK-STAT non-autonomously.

Lastly, we characterized peroxisome morphology and distribution in various tissues of adult female flies. Peroxisome is marked by PMP70, a peroxisomal membrane protein involved in the transport of long-chain acyl-CoA across peroxisomal membrane ^42^. We noticed that peroxisome showed tissue-specific enrichment in adult *Drosophila*. The peroxisome abundance was the lowest in the heart and highest in oenocytes, while the number of peroxisome in fat body (adipose-like) and pericardial cells (podocyte-like) was in the middle range (Figures 4h-4i). High enrichment of peroxisome is a key feature of mammalian liver ^14^. Interestingly, unlike oenocyte-specific manipulation, knocking down *Pex5* in fat body and midgut using *S106^GS^-Gal4* did not induce cardiac arrhythmia (Figure 4j). These results suggest that oenocyte-specific peroxisomal import contributes significantly to the over-production of pro-inflammatory cytokine and non-autonomous regulation of cardiac function. Here, we refer to the cellular stress responses (e.g., production of inflammatory cytokines) caused by impaired peroxisomal import as peroxisomal import stress (PIS).

### Pex5 KD-mediated peroxisomal import stress (PIS) up-regulates *Upd3* through JNK signaling

We next examined the molecular mechanism by which *Pex5* KD-mediated peroxisomal import stress (PIS) induces *Upd3* expression. Upd3 is known to play an important role in activating innate immunity and tissue repair upon infection. A recent genetic screening identified several transcription factors as the key regulators for infection-induced *Upd3* transcription, such as Mothers against dpp (Mad), the AP-1 complex (Kayak/Kay and Jun-related antigen/Jra), and Yorkie (Yki) ^43^. Interestingly, we found that oenocyte-specific knockdown of *Pex5* induced the expression of *Kay*, but not *Mad* and *Yki* (Figure 5a), suggesting Kay may be the transcription factor regulating *Upd3* expression upon PIS. Kay is the transcription factor downstream of JNK signaling ^44, 45^. We then asked whether PIS can also activate JNK signaling. Through immunostaining, we found that *Pex5* KD induced the phosphorylation of JNK in oenocytes (Figures 5b-5c). We further confirmed the results by measuring the expression of *Puckered* (*Puc*), the downstream target gene of Kay. Consistently, *Puc* expression was also induced by oenocyte-specific *Pex5* KD (Figure 5d). Interestingly, the activation of JNK pathway, indicated by the up-regulation of *Kay* and *Puc*, was also observed in flies with oenocyte-specific *Cat* KD, but not *ND75* KD (Figure 5e). This further suggests that peroxisome dysfunction plays a key role in activating JNK pathway. Finally, to directly examine the role of Kay in mediating PIS-induced *Upd3* transcription, we generated fly lines with *Pex5; Kay* double knockdown. The knockdown efficiency of *Kay* RNAi was verified by QRT-PCR (Figure S4e). As expected, knockdown of *Kay* completely blocked *Pex5* KD-induced *Upd3* transcription in oenocytes (Figure 5f), suggesting that *Pex5* KD-mediated PIS up-regulates *Upd3* through JNK signaling.

**Figure 5:**
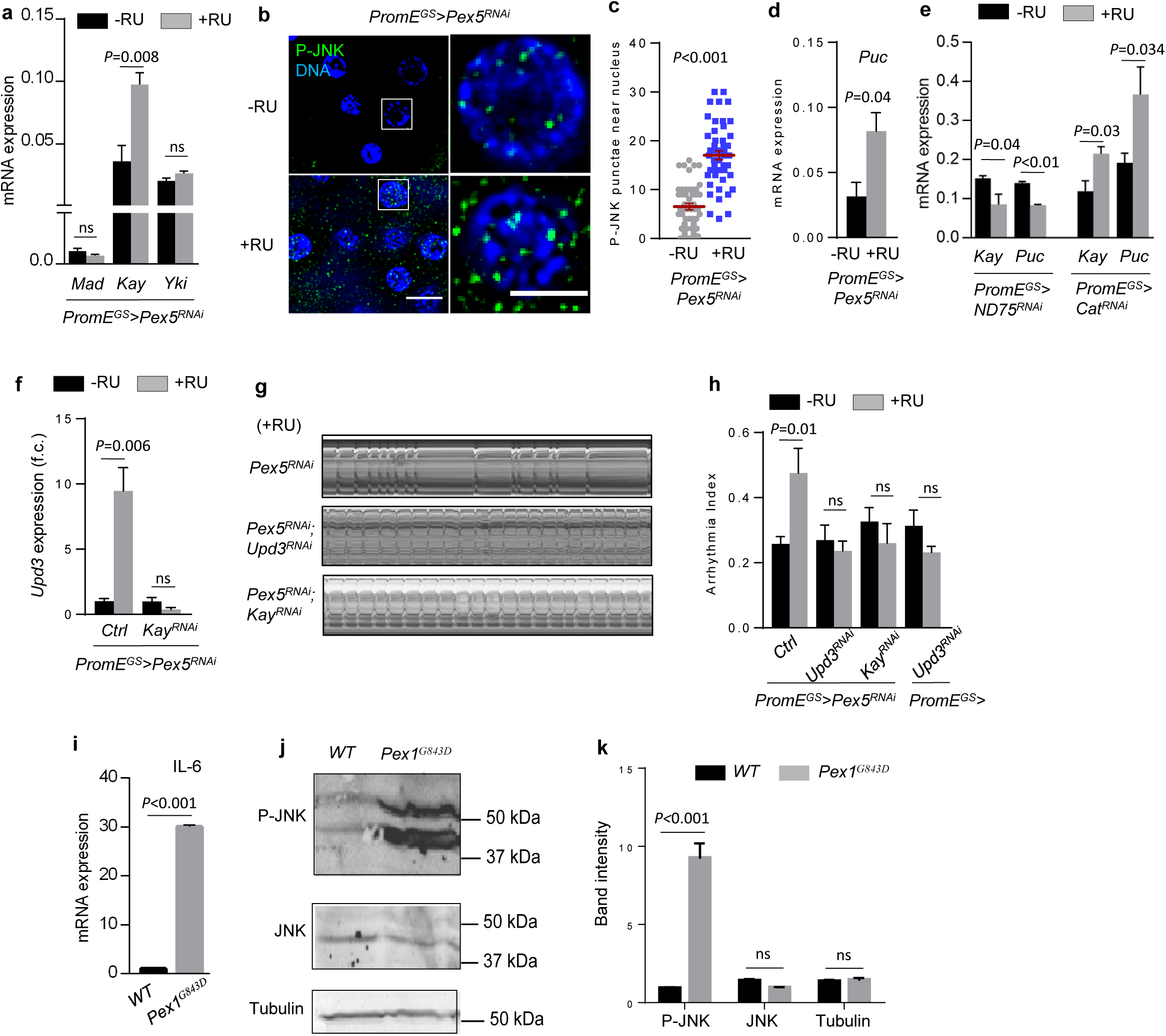
Pex5 KD-mediated peroxisomal import stress (PIS) upregulates *Upd3* through JNK signaling. **(a)** QRT-PCR analysis showing relative mRNA expression of *Mad*, *Kay*, *Yki* in oenocytes dissected from *PromE^GS^>Pex5^RNAi^* flies. **(b)** Representative images of P-JNK immunostaining of dissected oenocytes with (+RU) or without (-RU) *Pex5* KD. White boxes indicate the regions shown in the right insets. Scale bar: 6.7 µm. **(c)** Quantification of P-JNK punctae near oenocyte nuclei. N = 6. Dot plot shows the quantifications of 6 biological replicates (two images per replicates, five nuclei per image were analyzed). **(d)** QRT-PCR analysis showing relative mRNA expression of *Puc* in oenocytes dissected from flies with oenocyte-specific knockdown of *Pex5*. N = 4. **(e)** QRT-PCR analysis showing relative mRNA expression of *Kay* and *Puc* in oenocytes dissected from flies with oenocyte-specific knockdown of *ND75* and *Cat*. N = 4. **(f)** Relative mRNA expression of *Upd3* in oenocytes dissected from flies with *Pex5* KD or *Pex5; Kay* double KD. N = 4. **(g)** Representative M-mode traces of flies with oenocyte-specific *Pex5* KD, *Pex5; Upd3* double KD, or *Pex5; Kay* double KD. **(h)** Arrhythmia index of flies with oenocyte-specific *Pex5* KD, *Pex5; Upd3* double KD, *Pex5; Kay* double KD, and *Upd3* KD. N = 25. **(i)** Relative mRNA level of pro-inflammatory cytokine *IL-6* in human PEX1-G843D-PTS1 cells. N = 3. **(j)** Western blots showing the levels of P-JNK and JNK in wild-type and human PEX1-G843D-PTS1 cells. **(k)** Quantification of the band intensity of western blots in panel (j). N = 3. Data are represented as mean ± SEM. *P* values are calculated using unpaired *t-*test, ns: not significant.

To determine whether Upd3 and JNK signaling are required for oenocyte PIS-induced cardiac dysfunction, we analyzed the genetic interaction between *Pex5* and *Upd3* (or *Pex5* and *Kay*) in non-autonomous regulation of cardiac arrhythmia. As shown in Figures 5g-5h, knockdown of either *Upd3* or *Kay* specifically in oenocytes blocked *Pex5* KD-induced cardiac arrhythmia. *Upd3* KD alone did not affect cardiac arrhythmicity (Figure 5h). Thus, these data confirm that Upd3 is the primary factor produced in response to oenocyte-specific PIS and JNK activation to mediate oenocyte-heart communication.

Patients with Zellweger syndrome, due to the mutations in peroxins (e.g., Pex1 and Pex5), often develop severe hepatic dysfunction and show elevated inflammation ^16^. To test whether peroxisomal dysfunction in patients with Zellweger syndrome also induces the production of pro-inflammatory cytokines, we measured the expression of *IL-6* and the phosphorylation of JNK in PEX1-G843D-PTS1 cell line. This cell line is a human fibroblast cell line that was isolated, transformed and immortalized from patients with PEX1-p.G843D allele. Similar to *Pex5* KD flies, PEX1-G843D-PTS1 cells showed significant induction of *IL-6* transcripts (Figure 5i) and elevated phospho-JNK levels (Figures 5j-5k). Together, our data suggest an evolutionarily conserved mechanism for PIS-mediated induction of pro-inflammatory cytokines and activation of JNK signaling.

### Peroxisomal import function is significantly disrupted in aged oenocytes

Previous studies by us and others show that genes involved in peroxisomal import are down-regulated during aging ^13, 17^, suggesting an age-dependent impairment of peroxisomal import function. To monitor peroxisomal import during oenocyte aging, we first performed immunostaining using an anti-SKL antibody that recognizes peroxisomal matrix proteins containing the peroxisome targeting sequence (PTS), a SKL tripeptide sequence. We found that the number of SKL-positive punctae were significantly reduced in aged oenocytes, suggesting that the import of endogenous peroxisomal matrix proteins was impaired at old age (Figures 6a-6b). However, the reduced anti-SKL immunostaining might be caused by the decreased expression of the peroxisomal matrix proteins during aging ^13, 17^. To address this confounding effect, we directly examined peroxisomal import using a peroxisomal reporter in which YFP is engineered with a C-terminal PTS sequence (YFP-PTS). We transiently induced YFP-PTS reporter expression in young and aged oenocytes using *PromE^GS^-Gal4* (with 1-day RU486 feeding). Consistent with the results from anti-SKL immunostaining, aged oenocytes showed less YFP-PTS punctae, which indicates that fewer YFP-PTS proteins were imported into peroxisomes in aged flies (Figures 6c-6e).

**Figure 6:**
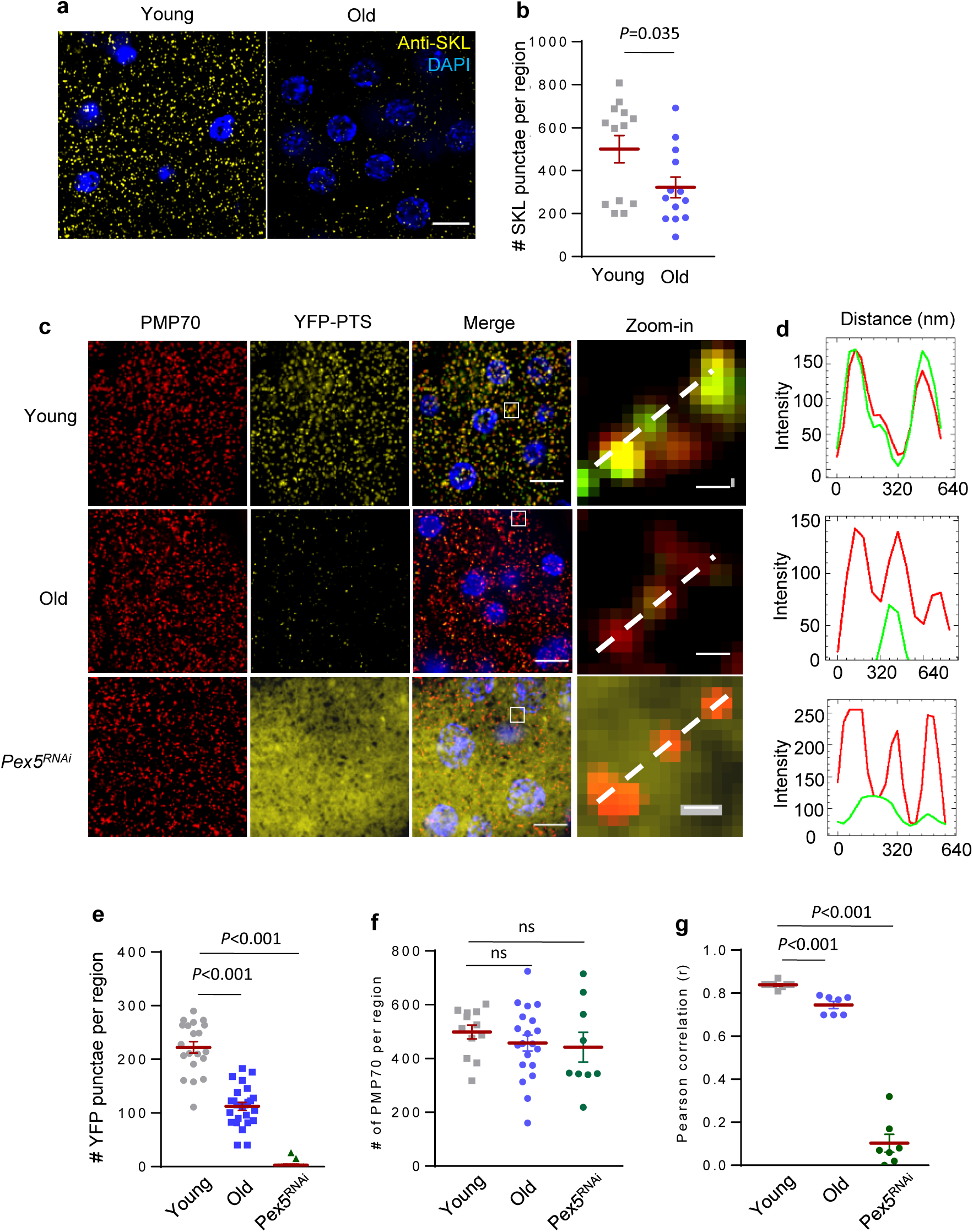
Peroxisomal import function is significantly disrupted in aged oenocytes. **(a)** Representative images of anti-SKL immunostaining of young and aged wild-type oenocytes. Scale bar: 6.7 µm. **(b)** Quantification on the number of SKL-positive punctae in Panel (a). N = 6. Dot plot shows the quantifications of 6 biological replicates, 2-3 ROIs per replicate. **(c)** Representative images to show co-localization of PMP70 and YFP-PTS in young and aged oenocytes, as well as oenocyte-specific Pex5 KD flies (*PromE>Pex5^RNAi^*) (Scale bar: 6.7 µm). Insets on the right show zoom-in peroxisome structures (the regions indicated by the white boxes in the merged panels, Scale bar: 281.6 nm). Hoechst 33342 was used for nuclear staining. **(d)** Line scan analysis to show the fluorescence intensity of PMP70 (red) and YFP-PTS (green) crossing peroxisomes in the insets (dashed line). **(e)** Quantification of the number of YFP-positive punctae in Panel (c). N = 6. Dot plot shows the quantifications of 6 biological replicates, 3-4 ROIs per replicate. **(f)** Quantification of the number of PMP70-positive punctae in Panel (c). N = 6. Dot plot shows the quantifications of 6 biological replicates, 3-4 ROIs per replicate. **(g)** Pearson correlation quantification measuring the correlation coefficiency of the co-localization between PMP70 and YFP-PTS in Panel (c). N = 6∼7. Data are represented as mean ± SEM. *P* values are calculated using unpaired *t-*test, ns: not significant.

To further confirm whether the YFP-PTS punctae are indeed localized with peroxisomes, we performed co-localization analysis between YFP-PTS and peroxisome marker PMP70. We found that the number of PMP70-positive peroxisomes did not change during oenocyte aging (Figures 6c, 6f). However, the co-localization of PMP70 and YFP-PTS was significant reduced in aged oenocytes according to the line scan analysis (Figure 6d) and Pearson’s correlation calculation (Figure 6g). Although the number of peroxisomes (PMP70-positive) did not change during aging, there were higher number of peroxisomes that showed either none or reduced YFP-PTS signals (Figures 6c-6d). The peroxisomes with no PTS-positive matrix proteins are known as “peroxisomal ghosts” (or non-functional peroxisomes), which is one of the cellular hallmarks of Zellweger Syndrome ^46, 47^. The decreased co-localization of PMP70 and YFP-PTS was also observed in oenocyte-specific *Pex5* KD flies (Figures 6c-6g). Together, these results suggest that peroxisomal import, but not peroxisomal biogenesis is impaired during oenocyte aging.

### Oenocyte-specific *Pex5* activation alleviates age-related PIS and preserves cardiac function

Although peroxisomal import function declines with age, the causal relationship between impaired peroxisomal import and tissue aging remains elusive. To determine the physiological significance of peroxisomal import in aging and cardiac health, we next asked whether preserving peroxisomal import function in oenocytes could attenuate the production of pro-inflammatory cytokines and protect hearts from paraquat- and aging-induced arrhythmicity. Our previous oenocyte translatomic profiling showed that Pex5 is down-regulated upon paraquat treatment and aging ^13^. We thus used a CRISPR/Cas9 transcriptional activation system ^48^ to drive physiologically relevant expression of *Pex5* from its endogenous locus to see if over-expression of *Pex5* in oenocytes could block age-related induction of *Upd3*, impairment of peroxisomal import, elevated cardiac arrhythmicity. We first combined a catalytically dead Cas9 line (*UAS-dCas9-VPR*) with oenocyte-specific driver line (*PromE^GS^-Gal4*), and then cross it with flies expressing *Pex5* guide RNA.

As expected, *Pex5* transcripts in oenocytes were induced about 2-fold after RU486 feeding (Figure 7a). Interestingly, *Pex5* over-expression attenuated the age-related *Upd3* induction in aged oenocytes (Figure 7b). *Pex5* over-expression also restored peroxisomal import function in aged oenocytes, as indicated by higher number of anti-SKL immunostaining (functional peroxisomes) (Figures 7c-7d) and lower percentage of peroxisomal ghosts (non-functional peroxisomes) (Figures 7c, 7e). *Pex5* over-expression did not change the total peroxisome numbers (PMP70-positive punctae) (Figures 7c, 7f). Our result is consistent with the previous study showing that over-expression of *Pex5p* partially restores the peroxisomal import function of *pex14Δ* mutants in yeast ^49^. Intriguingly, oenocyte-specific over-expression of *Pex5* blocked paraquat-induced arrhythmicity (Figures 7g-7h), and preserved cardiac function at old ages (Figures 7i-7j). Besides protecting cardiac function, oenocyte-specific over-expression of *Pex5* alleviated age-related ROS production in oenocytes (Figures 7k-7l). Together, our data show that preserving peroxisomal import function via *Pex5* over-expression can block the production of ROS and pro-inflammatory cytokine *Upd3*, and protecting hearts from stress- and aging-induced cardiomyopathy (Figure 7m). Thus, our studies provide strong evidence suggesting that age-related impairment of peroxisomal import is a novel and understudied cause of tissue aging.

**Figure 7:**
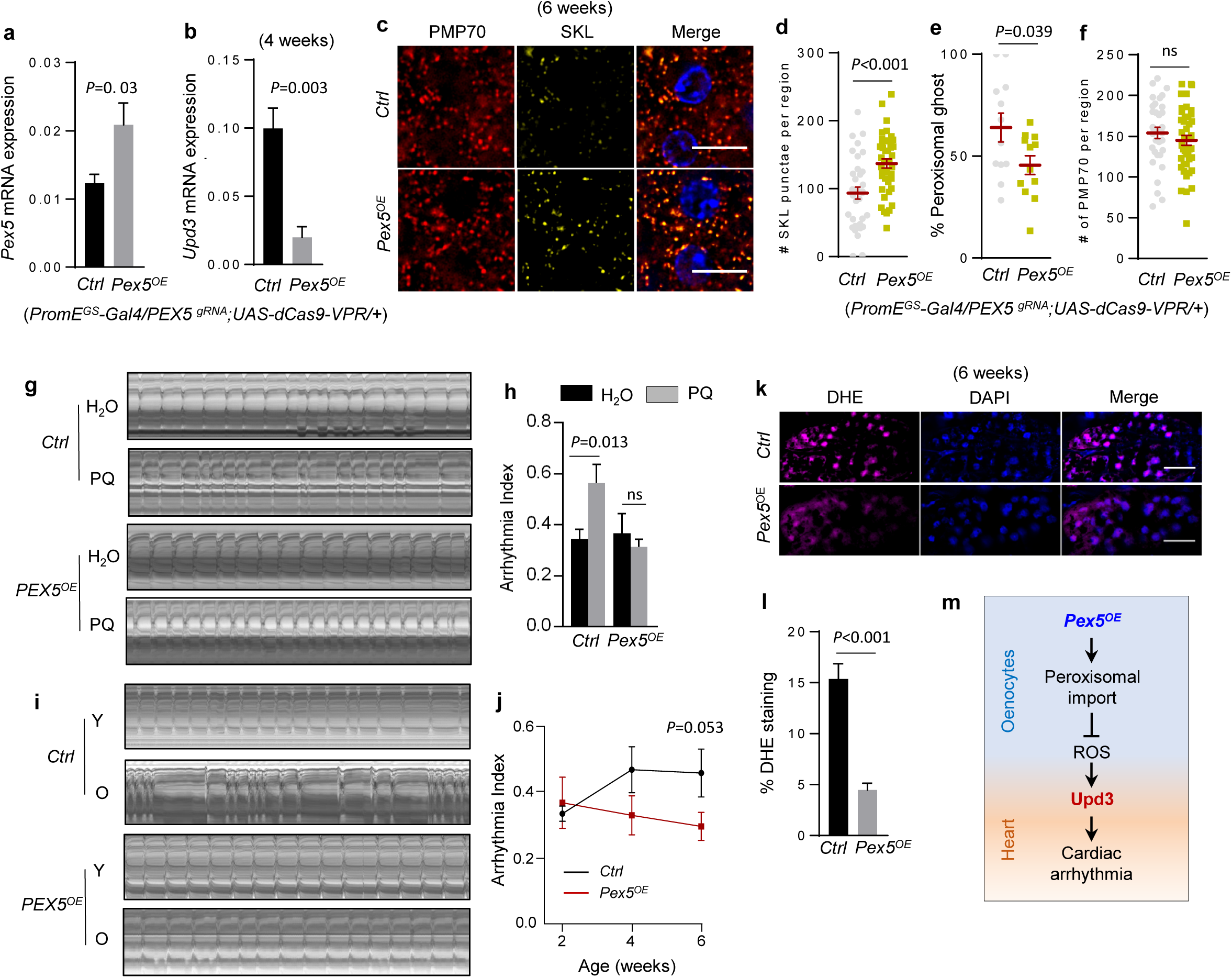
Oenocyte-specific *Pex5* activation alleviates age-related PIS and preserves cardiac function. **(a)** QRT-PCR analysis showing relative mRNA expression of *Pex5* in oenocytes dissected from flies with oenocyte-specific over-expression of *Pex5*. N=4. **(b)** QRT-PCR analysis showing relative mRNA expression of *Upd3* in 4-week-old oenocytes dissected from flies with oenocyte-specific over-expression of *Pex5*. N=4. **(c)** Representative images to show co-localization of anti-PMP70 and anti-SKL immunostaining in 6-week-old oenocytes dissected from flies with oenocyte-specific over-expression of *Pex5*. Hoechst 33342 was used for nuclear staining. Scale bar: 6.7 µm. **(d)** Quantification of SKL-positive punctae in Panel (c). N = 6. Dot plot shows the quantifications of 6 biological replicates, 3-4 ROIs per replicate. **(e)** Quantification of the percentage of peroxisomal ghosts (PMP70-punctae with no SKL signals) per region of interest (166.41 µm^2^) in Panel (c). N = 6. Dot plot shows the quantifications of 6 biological replicates, 2 ROIs per replicate. **(f)** Quantification of PMP70-positive punctae in Panel (c). N = 6. Dot plot shows the quantifications of 6 biological replicates, 3-4 ROIs per replicate. **(g)** Representative M-mode traces of flies with oenocyte-specific *Pex5* over-expression under paraquat treatment. **(h)** Arrhythmia index of flies with oenocyte-specific *Pex5* over-expression under paraquat treatment. N = 15. **(i)** Representative M-mode traces of flies with oenocyte-specific *Pex5* over-expression during normal aging. (J) Arrhythmia index of flies with oenocyte-specific *Pex5* over-expression during normal aging (2-week, 4-week and 6-week-old). N = 15. **(k)** DHE staining in 6-week-old oenocytes from flies with oenocyte-specific over-expression of *Pex5*. Scale bar: 20 µm. **(l)** Quantification of the percentage of DHE-positive staining in a Panel (k). N = 6. **(m)** Proposed model to show oenocyte Pex5-regulated peroxisomal import, ROS, Upd3 production, and non-autonomous regulation of cardiac function. Data are represented as mean ± SEM. *P* values are calculated using unpaired *t-*test, ns: not significant.

## Discussion

Here our studies provide direct evidence linking oenocyte/liver dysfunction to cardiac health. We also demonstrate that impaired peroxisomal import at old ages is the root causes of age-related chronic and systemic inflammation (inflammaging), whereas maintaining peroxisomal import slows tissue aging. Our genetic analyses show that aging-induced production of pro-inflammatory cytokine Upd3 from oenocytes/liver, in response to impaired hepatic peroxisomal import, activates JAK-STAT signaling in the heart to promote cardiac arrhythmia. Interestingly, either blocking *Upd3* production or over-expressing *Pex5* in oenocytes alleviates aging- or paraquat-induced cardiac dysfunction. Together, our studies established the vital role of hepatic peroxisomal import in the activation of systemic inflammation and non-autonomous regulation of cardiac aging. Our findings suggest that protecting oenocyte/liver function, in particular the peroxisomal import, is critical for prolonging cardiac healthspan.

IL-6 is the most important pro-inflammatory cytokine that is associated with inflammaging and age-related diseases ^5^. The up-regulation of IL-6 is often seen upon tissue injury, such as ischemia-reperfusion ^50^. IL-6 is also found to be secreted as a myokine from skeletal muscle during exercise ^51^, which is thought to mediate the beneficial effects of exercise. However, the studies using IL-6 knockout mouse models suggest that IL-6 is pathogenic and pro-inflammatory. For example, IL-6-deficient mice show reduce expansion of CD4^+^ T cells and autoimmune response ^52^. In *Drosophila*, IL-6-like cytokine Upd3 plays a key role in intestinal stem cell homeostasis ^53–55^, reproductive aging ^56^, glucose homeostasis and lifespan ^29^. Although unpaired family members, especially Upd2, have been shown to relays nutrient signals from fat body to brain ^57^, the role of Upd3 in inter-tissue communication and systemic aging regulation has not been fully established. For example, it is not known which tissue is the main source of Upd3 production, and which tissues (and tissue function) Upd3 targets during normal aging. In the present study, we show that Upd3, but not Upd1 and Upd2, mediates the communication between oenocytes and hearts. Among all tissues tested, oenocytes produce highest amount of Upd3 at old ages. Interestingly, only oenocyte-produced Upd3, not gut/fat body, modulates cardiac function during aging. Thus, our studies demonstrate that oenocytes are the main source of pro-inflammatory cytokine Upd3 in aged animals, and oenocytes may act as a hub for systemic aging control. Although it remains to be determined why Upd3 produced from different tissues exhibits distinct roles, it will be interesting to test whether oenocyte-produced Upd3 can target tissues beyond the heart (e.g., the central nervous system).

It is well-known that mitochondria play an important role in regulating chronic inflammation (inflammaging) ^5^. There is a strong interplay between mitochondria and peroxisomes in the maintenance of intracellular redox homeostasis ^58^. In fact, peroxisome deficiency can alter mitochondrial morphology and respiration ^59, 60^, and induce mitochondria-mediated cell death ^61^. However, the role of peroxisome in inflammaging still remains elusive. In the present study, we discover that knockdown of peroxisomal import receptor *Pex5* triggers a significant production of pro-inflammatory cytokine Upd3 in a JNK-dependent manner. Although it remains to be solved how impaired peroxisomal import activates JNK signaling pathway, it is clear that prolonged production of Upd3 from oenocytes is detrimental to cardiac function under aging. Interestingly, peroxisomal import stress (PIS) in both flies and human fibroblasts of Zellweger spectrum patients activates JNK pathway and the production of IL-6-type cytokines. These findings suggest a conserved cellular response to impaired peroxisomal import. Upd3/IL-6 might represent a group of inflammatory factors (we name them as “peroxikines”) that response specifically to peroxisome import stress. These factors are essential in maintaining tissue homeostasis upon PIS, since we can block the production of these factors to alleviate the detrimental effects of impaired peroxisomal import. The expression of the peroxikinies can also serve as biomarkers for cellular peroxisomal import stress and potentially inflammaging.

Age-related impairment of peroxisomal import has been previously reported in human senescence cells and nematodes ^17–19^, although the underlying mechanism is poorly understood. One hypothesis is that Pex5 protein is extensively oxidized during aging, which leads to the polyubiquitination and degradation of Pex5 (Ma et al., 2013). In senescent human fibroblasts, Pex5 is found to accumulate on the peroxisomal membrane (Legakis et al., 2002), suggesting that Pex5 recycling might be impaired during senescence. The dysregulation of Pex5 recycling could be caused by either reduced activities of ubiquitin ligases (Pex2 and Pex12). Or it is due to age-dependent decreases in ATP production, as Pex5 recycling is an ATP-dependent process ^62^. Finally, the impaired peroxisomal import could be simply due to the decreased expression of *Pex5* at old ages, as we observe a two-fold reduction of actively translated *Pex5* mRNA in aged oenocytes ^13^. Therefore, the amount of functional Pex5 are significantly reduced during normal aging, which explains why over-expressing *Pex5* can restore the import function and delay age-related pathologies. It should be noticed that excess expression of *Pex5* can block the normal import function ^63^. We utilized a CRISPR/Cas9 activation system to achieve optimal expression of endogenous *Pex5*, which has proven to be effective way to restore impaired peroxisomal import function and attenuate aging-induced cardiac dysfunction. Thus, our genetic analysis of Pex5 establish peroxisomal import as a novel root cause of tissue aging.

Besides the regulation of redox homeostasis, peroxisome performs several other essential functions, such as fatty acid beta-oxidation and ether phospholipid biosynthesis. Therefore, besides redox imbalance, impaired peroxisomal import could also lead to the accumulation of very long chain fatty acids and reduced production of anti-inflammatory lipids, such as docosahexaenoic acids (DHA) ^64^. DHA is known as the precursor of a group of anti-inflammatory lipids, such as resolvins, maresins and protectins. Thus, the decreased of DHA production, due to impaired peroxisomal import, can promote chronic inflammation in the liver and induce cardiomyopathy at old ages.

In summary, our studies reveal that *Drosophila* oenocytes (hepatocyte-like tissue) play a vital role in non-autonomous regulation of cardiac aging. Oenocytes produce pro-inflammatory cytokine Upd3 in response to peroxisomal import stress to modulate heart function. Protecting oenocytes from PIS and Upd3 production can alleviate cardiomyopathy under oxidative stress or aging. Our findings suggest that peroxisome is a vital organelle and central regulator of inflammaging. Maintaining healthy peroxisome import function (especially in oenocytes/liver) offers a promising strategy to combat age-related cardiac diseases.

## Methods

Detailed reagent information is provided in Supplemental Table S1.

### *Drosophila* husbandry and strains

A detailed list of fly strains is provided in Supplemental Table S1. The following genotypes were used as control in the knockdown or over-expression experiments: *w^1118^*, *GAL4^RNAi^* (BDSC#35783), *y^1^ v^1^; P[CaryP]attP2* (BDSC #36303), or *y^1^ v^1^; P[CaryP]attP40* (BDSC # 36304). Two independent RNAi lines were used in the knockdown of *Upd3*, *Pex5*, and *ND75*. Female flies were used in all experiments. Flies were maintained at 25°C, 60% relative humidity and 12-hour light/dark cycle. Adults and larvae were reared on a standard cornmeal and yeast-based diet, unless otherwise noted. The standard cornmeal diet consists of the following materials: 0.8% cornmeal, 10% sugar, and 2.5% yeast. RU486 (mifepristone, Fisher Scientific) was dissolved in 95% ethanol, and added to standard food at a final concentration of 100 µM for all the experiments except for *Pex5* over-expression, where 200 µM of RU486 was used. For paraquat feeding assay, paraquat (dichloride hydrate pestanal, Sigma) was dissolved in distilled water to a final concentration of 10 mM (Figures 1-3) or 20 mM (Figures 4-7). About 150 µl of paraquat working solution was prepared freshly and added onto the surface of fly food. Paraquat solution was air dried for 20 minutes at room temperature prior to use.

### Human PEX1-G843D-PTS1 cell culture

The human PEX1-G843D-PTS1 cell lines were from the Braverman laboratory ^65^. It is a fibroblast cell line that was originally isolated, transformed and immortalized from patients with PEX1-p.G843D, a missense allele that accounts for one-third of all Zellweger spectrum disorder alleles. It stably expresses GFP-peroxisome targeting signal 1 (PTS1) reporter. The PEX1-G843D-PTS1 cell lines were cultured in DMEM with 10% FBS at 5% CO_2_. The wild-type cell lines were established from the fibroblasts of healthy donors.

### Fly heartbeat analysis

Cardiac contractility was measured using semi-intact female *Drosophila* hearts as previously described ^66^. High speed movies (30s, 100 frames/second) were recorded using a Hamamatsu ORCA-Flash 4.0 digital CMOS camera (Hamamatsu) on an Olympus BX51WI microscope with a 10X water immersion lens (Olympus). The live images taken from the heart tub within abdominal A3 segment were processed using HCI imaging software. M-modes and cardiac parameters were generated using SOHA, a MATLAB-based image application ^66^. A fifteen-second of representative M-mode was presented in the figures to show the snapshot of the movement of heart wall over time. Arrhythmia index (AI) is calculated as the standard deviation of all heart periods in each fly normalized to the median heart period. Heart period is the pause time between the two consecutive diastole ^66^. At least 15 fly hearts were dissected and analyzed for each group in all the experiments.

### DHE staining

ROS detection was performed using DHE dye (dihydroethidium, Fisher Scientific) following a previously published method ^67^, with minor modification. Female oenocytes were dissected in 1x PBS according to a published protocol ^68^. Abdominal fat body was removed by liposuction and fly carcass with intact oenocytes was incubated in 30 μM of DHE solution for 5 minutes in the dark. After washing with PBS three times, the oenocytes were staining with Hoechst 33342 (1 μg/ml) (ImmunoChemistry Technologies) for 10 minutes, mounted in ProLong Gold antifade reagent (Thermo Fisher Scientific), and imaged with an epifluorescence-equipped BX51WI microscope (Olympus).

### Secretory factor screen

*Drosophila* secretory proteins were first identified from the Gene List Annotation for *Drosophila* (GLAD) ^28^ and compared to our recent oenocyte translatomic analysis ^13^. Candidate genes that are differentially expressed in aged or PQ-treated oenocytes were selected in a RNAi screening using oenocyte-specific driver *PromE-Gal4*. About 20 mated females (3∼5-day-old) per genotype were treated with 10 mM paraquat (as described above) for 24 hours before heart beat analysis (SOHA).

### RNA extraction and quantitative RT-PCR

Adult tissues (oenocyte, heart, gut, fat body, pericardial cells) were dissected according to previously published protocols ^68, 69^. For oenocyte dissection, we first removed fat body through liposuction and then detached oenocytes from the cuticle using a small glass needle. Tissue lysis, RNA extraction, and cDNA synthesis were performed using Cells-to-CT kit (Thermo Scientific).

Quantitative RT-PCR (QRT-PCR) was performed with a Quantstudio 3 Real-Time PCR system and PowerUp SYBR Green Master Mix (Thermo Fisher Scientific). Two to three independent biological replicates were performed with two technical replicates. The mRNA abundance of each candidate gene was normalized to the expression of *RpL32* for fly samples and *GAPDH* for human samples, by the comparative C_T_ methods. Primer sequences are listed in the Supplemental Table S2.

### Antibody and immunostaining

The following antibodies were used in immunostaining: anti-GFP (Cell Signaling Technology, #2956S, 1:200), anti-P-JNK (Thr183/Tyr185) (Cell Signaling Technology, #4668S, 1: 100), anti-STAT (1:900, a gift from Steven X. Hou), anti-PMP70 rabbit polyclonal antibody (1:200, generated against the *Drosophila* PMP70 C-terminal region 646-665 (DGRGSYEFATIDQDKDHFGS) by Pacific Immunology), and anti-SKL rabbit polyclonal antibody (1:250, raised by Richard Rachubinski). An anti-PMP70 Guinea Pig polyclonal antibody (A gift from Kyu-Sun Lee, 1:500) was used in peroxisome import assay (Figure 7). Secondary antibodies were obtained from Jackson ImmunoResearch.

Adult tissues were dissected and fixed in 4% paraformaldehyde for 15 min at room temperature. Tissues were washed with 1x PBS with 0.3% Triton X-100 (PBST) for three times (∼5 minutes each time), and blocked in PBST with 5% normal goat serum (NGS) for 30 minutes. Tissues were then incubated overnight at 4°C with primary antibodies diluted in PBST, followed by the incubation with secondary antibodies for two hours at room temperature. After wash, tissues were mounted using ProLong Gold antifade reagent (Thermo Fisher Scientific) and imaged with an epifluorescence-equipped BX51WI microscope (Olympus). Hoechst 33342 was used for nuclear staining.

### Image analysis and quantification

Fluorescence images were first processed and deconvoluted using Olympus CellSens Dimensions software (Olympus). The number of puncta or fluorescent intensity/area in a selected region of interest (ROI, 918.33 µm^2^ for Figures 5-6, 367.29 µm^2^ for Figure 4&7) was measured using the CellSens “Measure and Count” module after adjusted the intensity threshold to remove background signals. Two to four ROIs were analyzed for each image. The imaging quantifications were done single or double blind.

Co-localization analysis was performed in Image J. Coloc 2 plugin function was used to calculate Pearson’s correlation (r). Line scan analysis in Figure 6 was performed by creating composite images containing SKL and PMP70 immunostaining. We then drew a line crossing three peroxisomes in a selected ROI, and intensity plots were generated using multi-channel plot profile in the BAR package (https://imagej.net/BAR).

To identify peroxisomal ghost, images were processed, and thresholds were adjusted following the procedures described above. Image segmentation was performed for each channel (SKL: green, PMP70: red) using Otsu thresholding. Then SKL and PMP70 channels were merged in ImageJ. For each merged image, three ROIs (166.41 µm^2^) were selected and the total number of peroxisome and peroxisomal ghosts (PMP70-positive punctae with no SKL signals) were manually counted. The percentage of peroxisomal ghosts was presented in the figures.

### Hemolymph extraction

Hemolymph was extracted by piercing into the thoraces of 30 adult female flies (each genotype) with glass needles made of capillary tubes using Sutter Puller (Sutter Instrument, Model P-97). The following settings were used to prepare glass needle: Heat = 302, Pull =75, Vel = 75, Time = 155, P = 435. Vacuum was used to facility the hemolymph extraction. About 0.5 µl of hemolymph was collected in a 1.5 ml microcentrifuge tube containing 1x PBS and protein inhibitor cocktail. After quick centrifuge at 3000 x RPM for 2 minutes at 4°C, the hemolymph samples were snap-freeze with liquid nitrogen and stored at −80°C for western blot analysis.

### Western blot analysis

Proteins samples were denatured with Laemmli sample buffer (Bio-Rad, Cat# 161-0737) at 95°C for 5 minutes. Then proteins were separated by Mini-PROTEAN® TGX Precast Gels (Bio-Rad). Following incubation with primary and secondary antibodies, the blots were visualized with Pierce ECL Western Blotting Substrate (Thermo Scientific). The following antibodies were used: anti-GFP (Cell Signaling Technology, #2956S, rabbit, 1:1000), anti-P-JNK (Cell Signaling Technology, #9255, 1:2000), anti-JNK (Cell Signaling Technology, #9252, 1:1000), anti-Tubulin (Sigma, #T5168, 1:2000). For hemolymph samples, 4–15% Mini-PROTEAN® TGX Stain-Free™ Precast Gels (Bio-Rad, Cat# 456-8085) was used. The Stain-Free^TM^ gel was UV activated (Bio-Rad) using ChemiDoc MP Imager to visualize total protein loading before immunoblotting.

### Statistical analysis

GraphPad Prism (GraphPad Software, version 6.07) was used for statistical analysis. All *p* values represent unpaired two-tailed Student’s t-test. In fly heartbeat analysis, the outliers were excluded using Robust regression and Outlier removal (ROUT) method (Q=1%) prior to the data analysis ^70^.

## Acknowledgements

We thank Bloomington Drosophila Stock Center and Harvard Medical School for fly stocks. We thank Steven X. Hou, Richard Rachubinski, and Kyu-Sun Lee for antibody reagents, Marc Tatar, Alex Gould, Doug Harrison, Heinrich Jasper, Rolf Bodmer, and Erika Bach for fly stocks. We thank Nancy Braverman for *Pex1* mutant cell lines. We also thank Rolf Bodmer and Karen Ocorr for help with fly heartbeat analysis.

## Author Contributions

KH and HB conceived the study and wrote the manuscript. KH performed most of the experiments. TM performed Upd3 secretion assay, KC performed SOHA analysis on cardiac JAK-STAT activation. PK performed tissue-specific Upd3 expression measurement. QJ performed part of the secretory factor screen. FD performed the analyses on Pex1 patient cells. AJS and FD guided the design of peroxisomal function analysis and generated PMP70 antibodies. KH, FD and HB analyzed the data. All authors reviewed and approved the manuscript.

---

**Figure S1.**
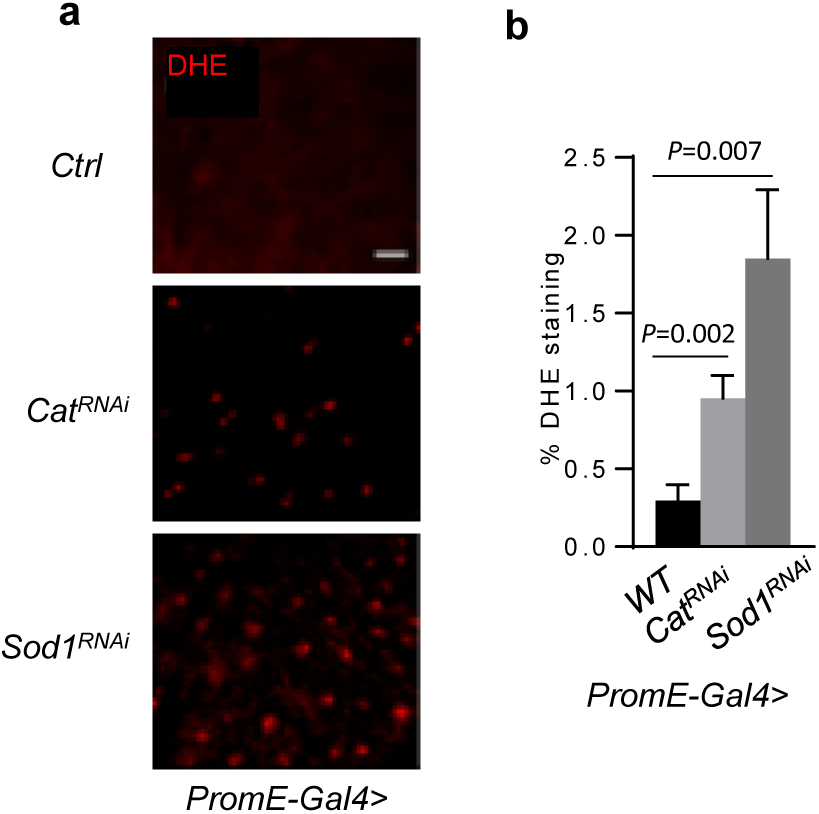
**(a)** Representative images to show DHE staining in oenocytes dissected from control and oenocyte-specific *Sod1* KD flies (*PromE-Gal4>UAS-Sod1^RNAi^*). Hoechst 33342 was used for nuclear staining. Scale bar: 20 µm **(b)** Quantification of relative DHE staining in control and oenocyte-specific *Sod1* KD flies (*PromE-Gal4>UAS-Cat^RNAi^*). Data are represented as mean ± SEM. *P* values are calculated using unpaired *t-*test, ns: not significant.

**Figure S2.**
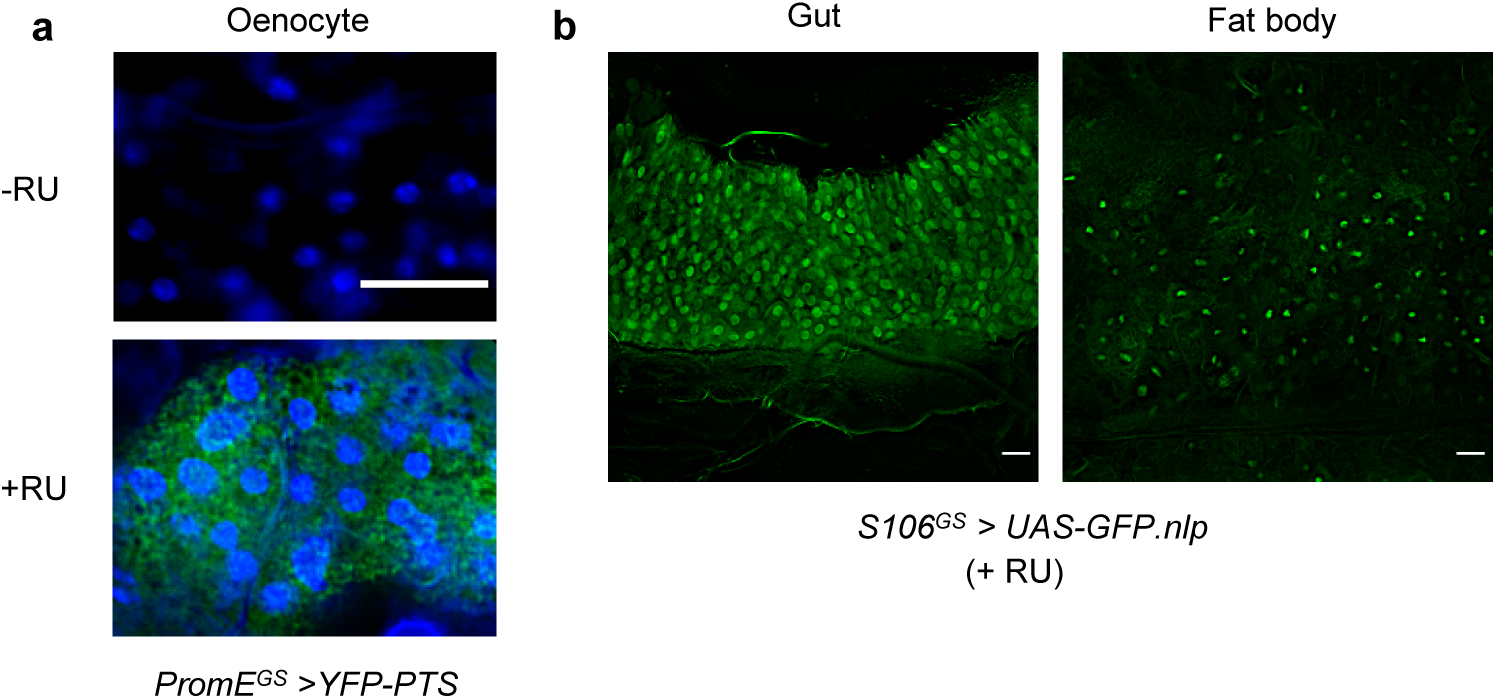
**(a)** Verification of oenocyte-specific GeneSwitch driver (*PromE^GS^-Gal4*). RU: mifepristone (RU486). **(b)** Verification of gut/fat body-specific GeneSwitch driver (S106*^GS^-Gal4*). Scale bar: 20 µm.

**Figure S3:**
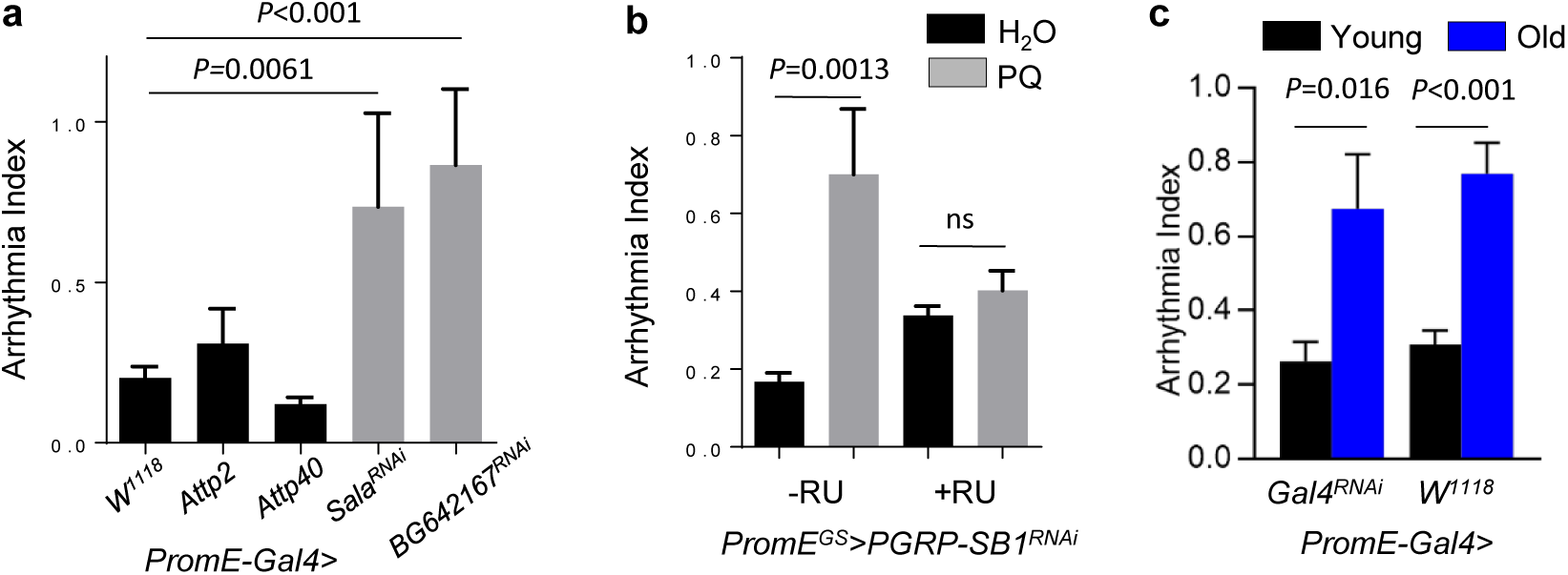
**(a)** Arrhythmia index of wild-type flies (*PromE-Gal4>w^1118^, PromE-Gal4>Attp2, PromE-Gal4>Attp40*) and oenocyte-specific knockdown of *Sala* and *BG642167*. N=12∼30. **(b)** Paraquat (PQ)-induced arrhythmia measured by SOHA for *PGRP-SB1* knockdown under oenocyte-specific GeneSwitch driver (*PromE^GS^-Gal4>PGRP-SB1^RNAi^*). N=15∼25. **(c)** Arrhythmia index of wild-type flies (*PromE-Gal4>UAS-Gal4^RNAi^* and *PromE-Gal4>w^1118^*) at young (2-week-old) and old age (6-week-old). N≥15. Data are represented as mean ± SEM. *P* values are calculated using unpaired *t-*test, ns: not significant.

**Figure S4.**
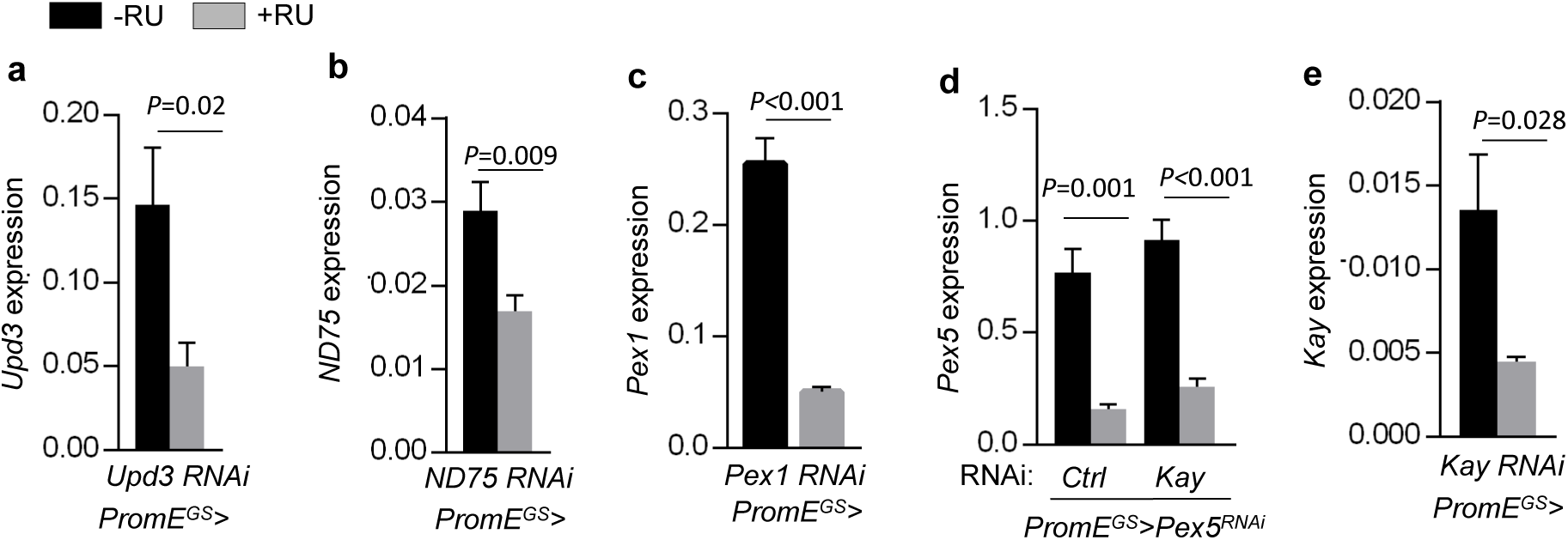
RNAi knockdown verification by QRT-PCR for *Upd3* **(a)**, *ND75* **(b)**, *Pex1* **(c)**, *Pex5* **(d)**, *Kay* **(e)**. Oenocyte-specific GeneSwitch driver (*PromE^GS^-Gal4*) was used. RU: mifepristone (RU486). Data are represented as mean ± SEM. *P* values are calculated using unpaired *t-*test, ns: not significant.

## Supplemental Tables

**Table S1:**
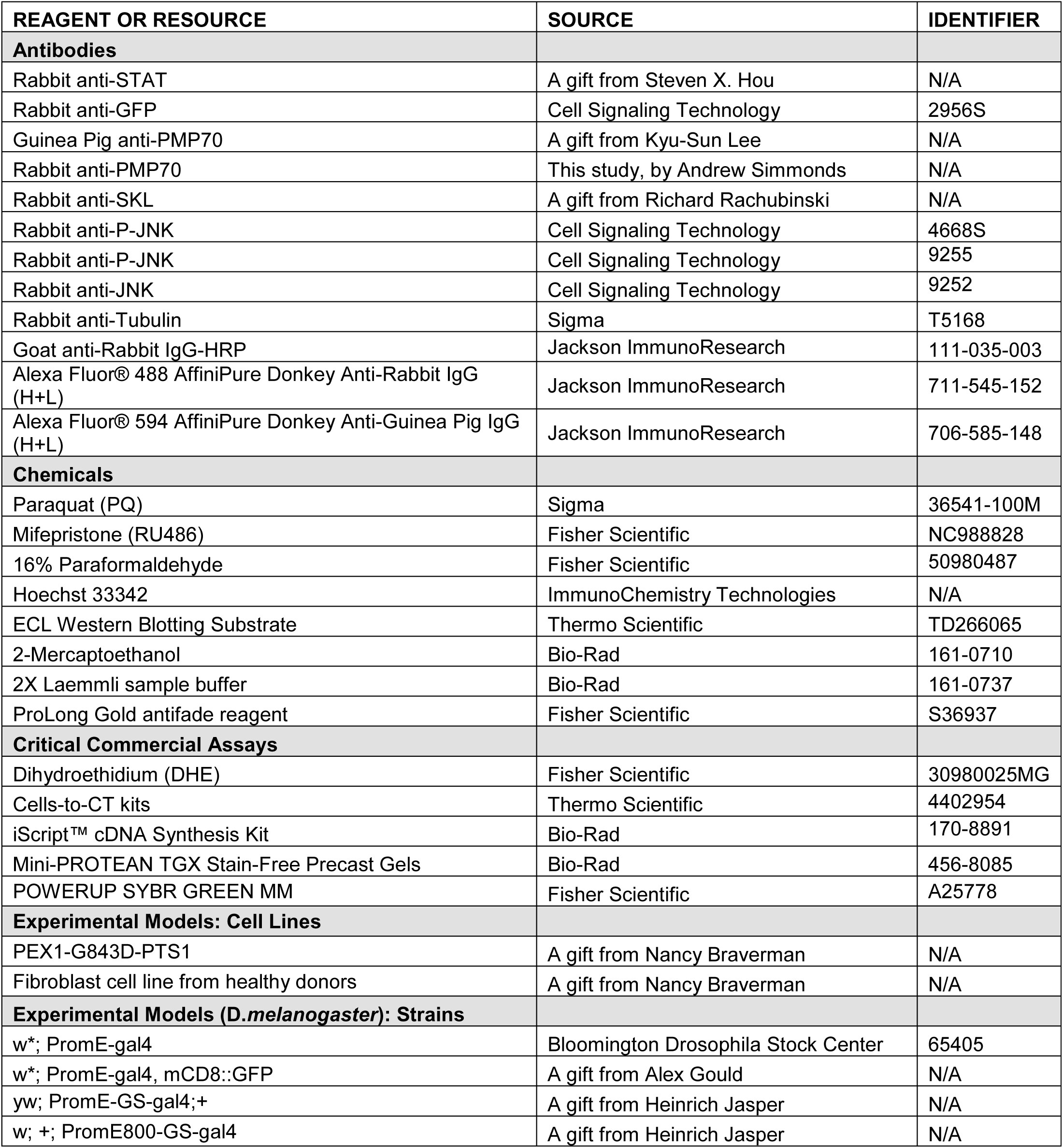
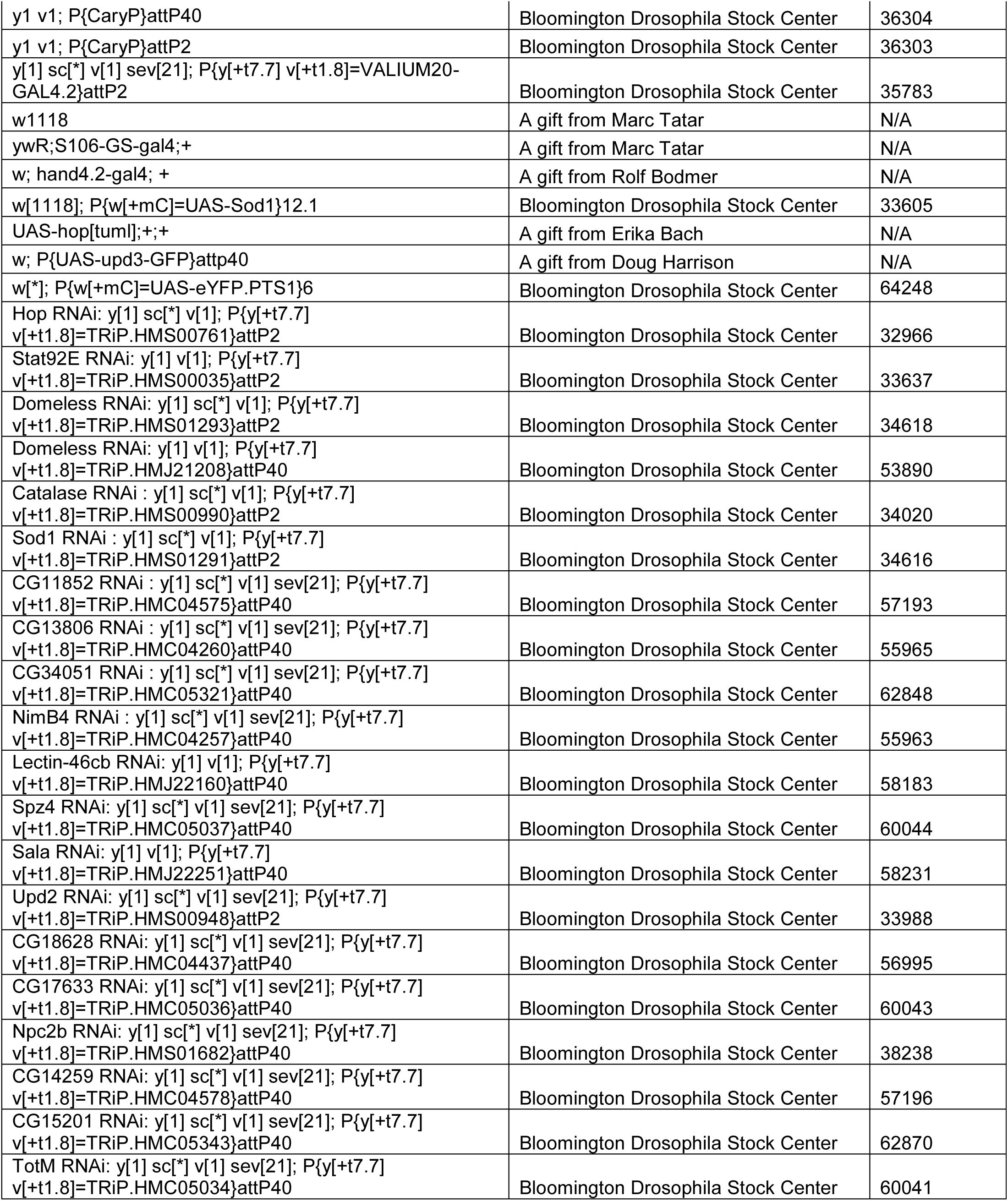
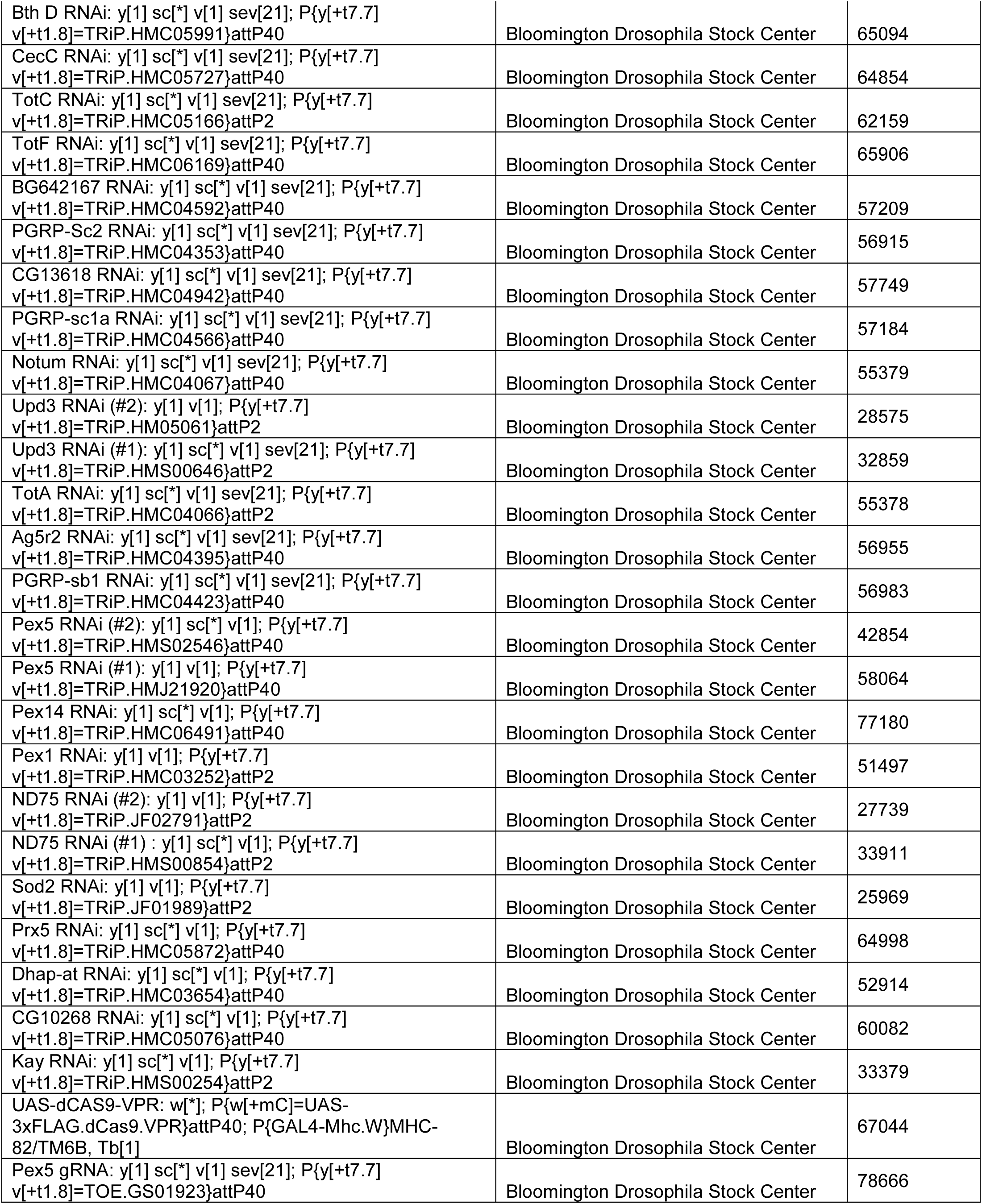
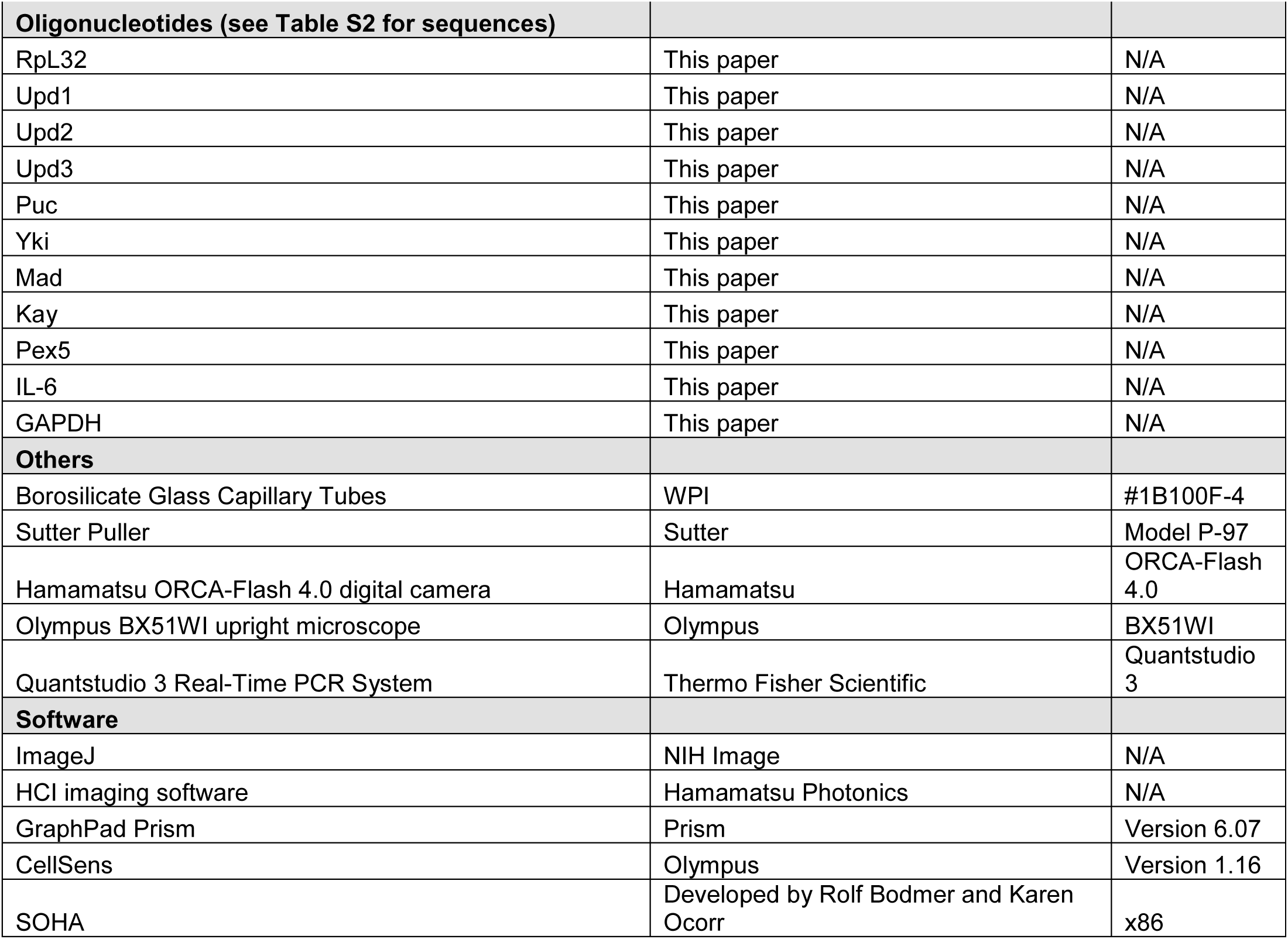
KEY RESOURCES TABLE.

**Table S2:**
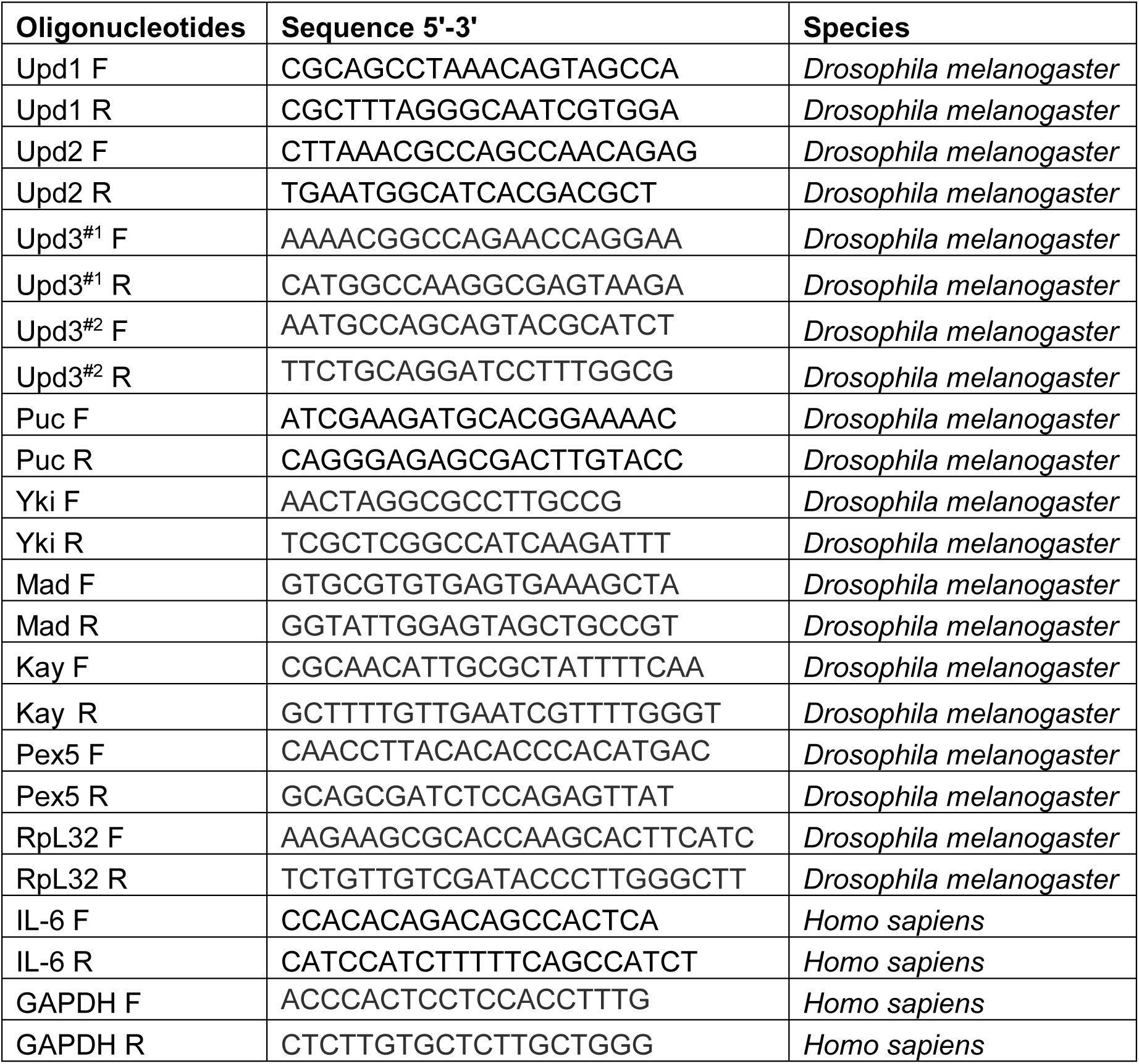
Primer list.

